# Structure of dimerized assimilatory NADPH-dependent sulfite reductase reveals the minimal interface for diflavin reductase binding

**DOI:** 10.1101/2024.06.14.599029

**Authors:** Behrouz Ghazi Esfahani, Nidhi Walia, Kasahun Neselu, Yashika Garg, Mahira Aragon, Isabel Askenasy, Hui Alex Wei, Joshua H. Mendez, M. Elizabeth Stroupe

**Author notes:** Corresponding Author: M. Elizabeth Stroupe Address: Florida State University 91 Chieftan Way Tallahassee, FL 32306.

## Abstract

*Escherichia coli* NADPH-dependent assimilatory sulfite reductase (SiR) reduces sulfite by six electrons to make sulfide for incorporation into sulfur-containing biomolecules. SiR has two subunits: an NADPH, FMN, and FAD-binding diflavin flavoprotein and a siroheme/Fe_4_S_4_ cluster-containing hemoprotein. The molecular interactions that govern subunit binding have been unknown since the discovery of SiR over 50 years ago because SiR is flexible, thus has been intransigent for traditional high-resolution structural analysis. We used a combination of the chameleon® plunging system with a fluorinated lipid to overcome the challenges of preserving a flexible molecule to determine a 2.78 Å-resolution cryo-EM structure of a minimal heterodimer complex. chameleon®, combined with the fluorinated lipid, overcame persistent denaturation at the air-water interface. Using a previously characterized minimal heterodimer reduced the heterogeneity of a structurally heterogeneous complex to a level that could be analyzed using multi-conformer cryo-EM image analysis algorithms. Here, we report the first near-atomic resolution structure of the flavoprotein/hemoprotein complex, revealing how they interact in a minimal interface. Further, we determined the structural elements that discriminate between pairing a hemoprotein with a diflavin reductase, as in the *E. coli* homolog, or a ferredoxin partner, as in maize (*Zea mays*).

**Significance Statement:** Sulfur is one of the essential building blocks of life. Sulfur exists in numerous redox states but only one can be incorporated into biomass – S^2-^ (sulfide). In *Escherichia coli*, a protein enzyme called sulfite reductase reduces sulfite by six electrons to make sulfide. Typical electron transfer reactions move one or two electrons at a time. The sequential transfer of two electrons three times to complete the conversion of sulfite to sulfide (or nitrite to ammonia) is unique to sulfite or nitrite reductases. *E. coli* SiR is a two-protein complex composed of a diflavin reductase flavoprotein and an iron metalloenzyme hemoprotein. Until now, the molecular interactions that govern subunit interactions remained a mystery because the extreme flexibility of the flavoprotein subunit, which has challenged X-ray or cryo-EM analysis for over 30 years. In overcoming these challenges, we used a combination of rapid plunging with a high critical-micelle-concentration lipid alongside a biochemically minimized complex to determine the 2.78 Å-resolution cryo-EM structure of a dimer between the flavoprotein and hemoprotein subunits.

---

Assimilatory sulfite reduction by NADPH-dependent sulfite reductase (SiR) is essential to produce sulfide for incorporation into sulfur-containing biomolecules. In γ-proteobacteria like *Escherichia coli*, SiR is a multimeric oxidoreductase composed of an octameric diflavin reductase (SiRFP) and four independently binding subunits of a siroheme-containing hemoprotein (SiRHP)^1–3^. Specifically, SiR catalyzes the six-electron reduction of sulfite (SO_3_^2-^), using three NADPH molecules that bind the SiRFP subunit. Each NADPH donates two electrons. The electrons first transfer to a SiRFP-bound FAD cofactor within an NADP^+^ ferredoxin reductase domain. The resulting FADH_2_ then transfers them to a SiRFP-bound FMN cofactor within a flavodoxin-like domain^4^. The electrons then transfer from the resulting FMNH_2_ cofactor to SiRHP, through a coupled siroheme-Fe_4_S_4_ cluster, and ultimately to the evolving substrate that binds to the active site siroheme iron^4^, which is housed in the SiRHP subunit, ultimately producing the fully reduced sulfide (S^2-^) product.

The *E. coli* SiRFP subunit is homologous to cytochrome P450 (CYP) reductase (CPR)^5^, the reductase domain of the bacterial CYP/CPR fusion CYP102A1/CYPBM3^6^, the reductase domain of nitric oxide synthase (NOSr)^7,8^, and methionine synthase reductase (MSR)^9^. One of the hallmarks of this diflavin reductase family is that they are exceptionally conformationally malleable because of a large conformational change of the two flavin binding domains that modulates electron transfer reaction amongst the NADPH, FAD, and FMN cofactors^10,11^. High resolution structural analysis is correspondingly challenging. For example, to date there are no high-resolution structures of the full-length NOS homodimer, the complex between methionine synthase and MSR, or the SiR heterododecameric holoenzyme (for simplicity, here referred to as a dodecamer). The structures of the CYP/CPR heterodimer and CYPBM3 are known. CYP/CPR form a 1:1 heterodimer^12–14^. The heme-binding and reductase domains are fused in CYPBM3^14^. Thus, little can be inferred about other homologs that function as higher-order protein complexes like SiR.

Like other well-studied diflavin reductases, SiRFP is highly modular (Fig. 1a and Tables 1 and S1). Despite its homology to other diflavin reductases, SiRFP is unique because it assembles into an octamer through its N-terminal 51 residues. Removing those residues results in a 60 kDa monomer (SiRFP-60), which binds SiRHP as a 1:1 heterodimer with reduced activity^3^. Further, removing the complete N-terminal FMN-binding flavodoxin (Fld) domain results in a 43 kDa monomer that contains just the NADPH- and FAD-binding NADP^+^ ferredoxin reductase (FNR) domain, along with an intervening connection domain (SiRFP-43), which also binds SiRHP as a 1:1 heterodimer but is inactive for electron transfer^15^. (Abbreviations and theoretical molecular weights are summarized in Tables 1 and S1).

**Figure 1:**
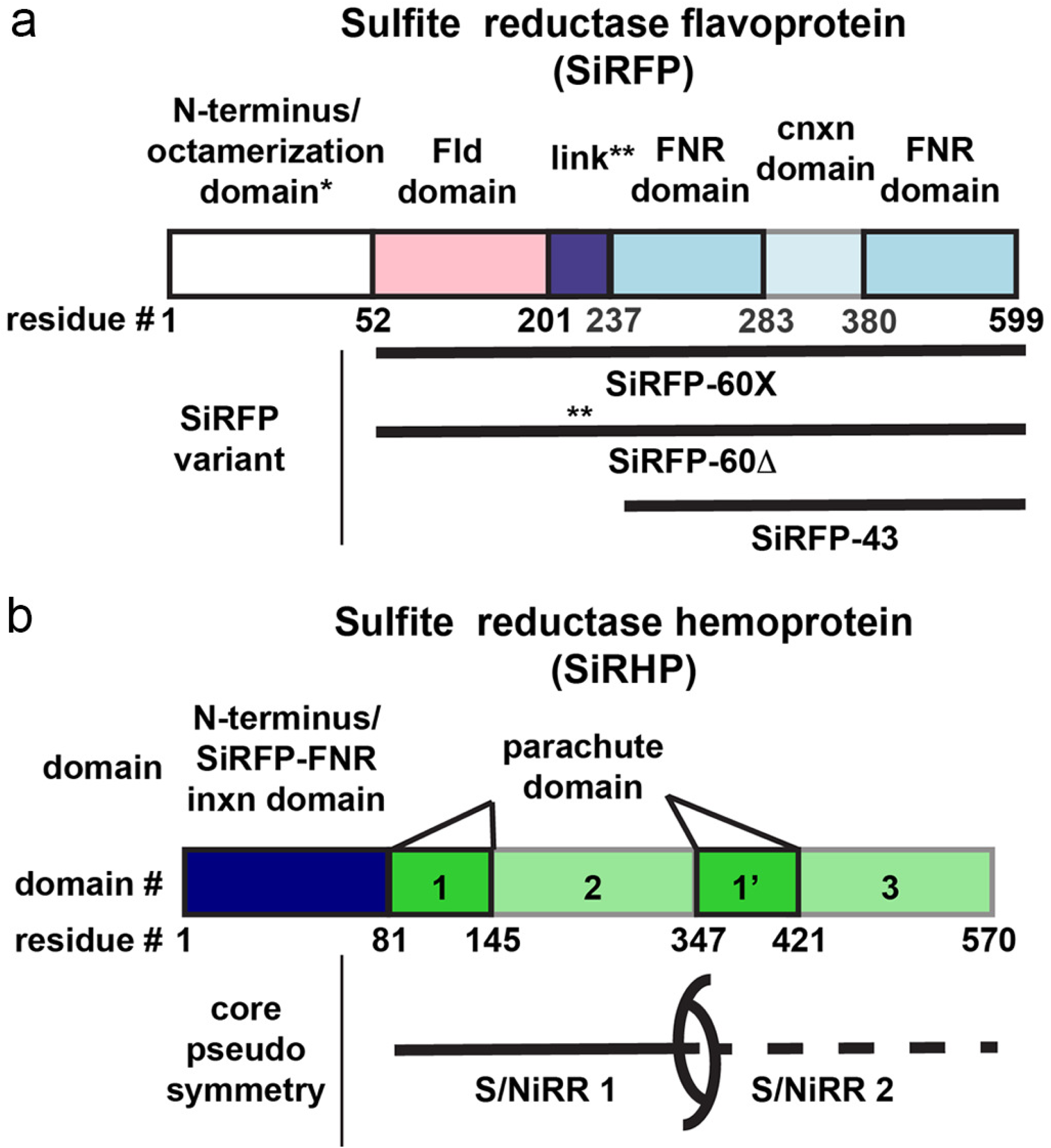
SiR components are highly modular. **a** SiRFP is composed of an N-terminal octamerization domain (not present in the constructs studied here), an FMN-binding Fld domain (pink) connected through a linker (purple) to an FAD-binding FNR domain (blue) interrupted by a connection (cnxn) domain (light blue). The variants and their names are denoted below: SiRFP-60X has a truncated octamerization domain (*) to make a 60 kDa monomer with engineered crosslinks between the Fld and FNR domain and a full-length linker. SiRFP-60Δ has a truncated octamerization domain (*) to make a 60 kDa monomer with a shortened linker (**, Δ212-217). SiRFP-43 only contains the FNR and connection domains to make a 43 kDa monomer. **b** SiRHP’s N-terminal 80 residues (dark blue) are solely responsible for forming a stable interaction with the FNR domain of SiRFP. The pseudosymmetric core is composed of sequential S/NiRRs (green and light green, domains 1/2 and 1’/3) that include a parachute domain (green, domains 1/1’).

**Table 1:**
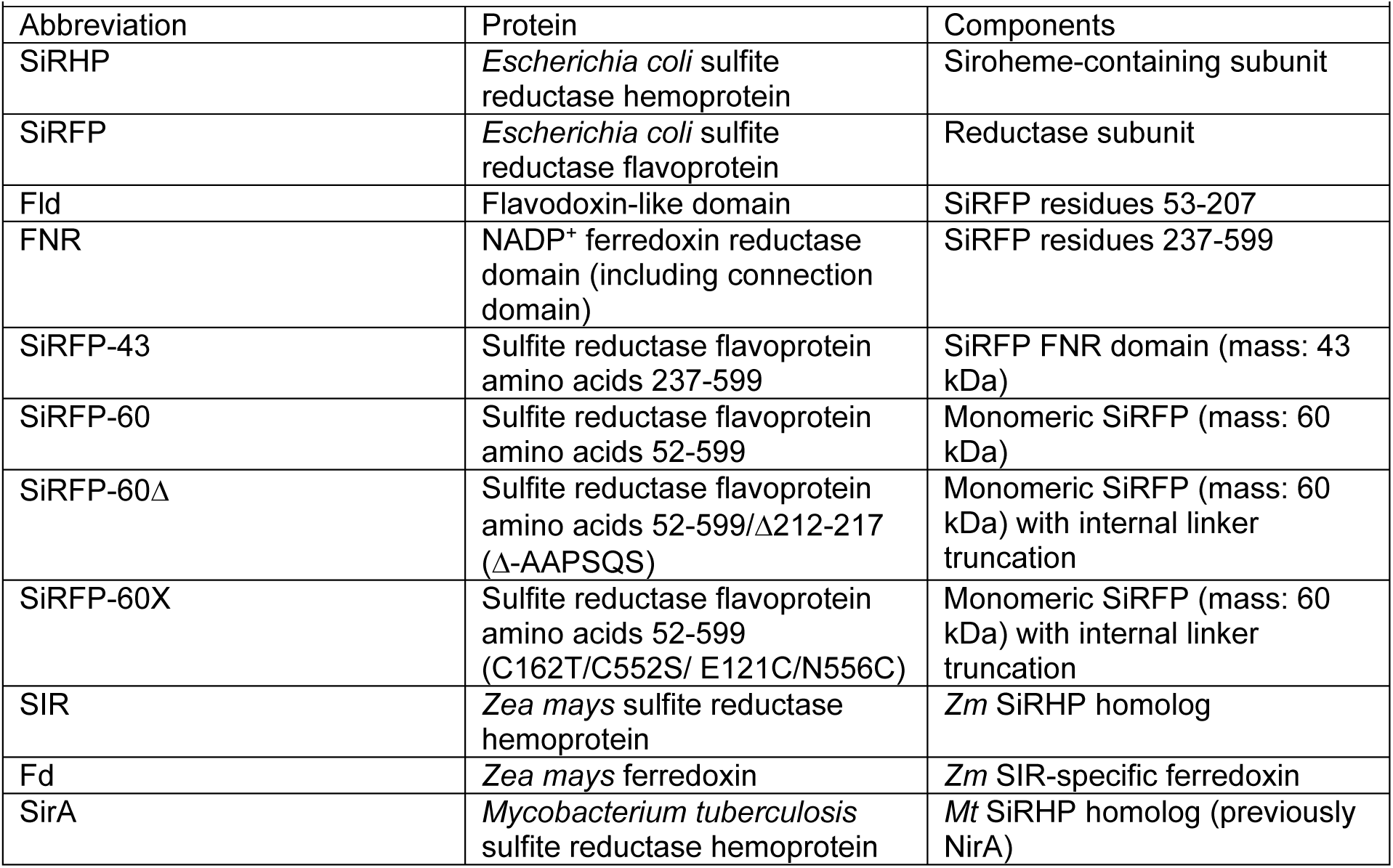
Sulfite Reductase Abbreviations.

SiRHP has few known homologs because of its unique siroheme/Fe_4_S_4_ cluster assembly that forms the sulfite-binding active site^16^. Siroheme is an iron-containing isobacteriochlorin that evolutionarily predates protoporphyrin IX-derived tetrapyrroles^17,18^. Assimilatory SiRs or siroheme-dependent nitrite reductases (NiRs) from other bacterial species or plants have a similar hemoprotein but use a transiently bound ferredoxin (Fd) as their electron source^19–21^. SiRs that are responsible for energy conversion, dissimilatory sulfite reductases (DSRs), share a common siroheme binding fold but are heterotetrameric and are often fused to auxiliary domains^22^. Their electron donors are poorly understood.

SiRHP houses the enzyme active site at the distal face of the siroheme iron^23–25^. Monomeric assimilatory SiRHP evolved through a gene-duplication, gene fusion event. Consequently, SiRHP has a pseudo-symmetric two-fold symmetry axis that runs through an α/β domain termed the parachute domain^23^ that relates two structurally similar sulfite/nitrite reductase repeats (S/NiRRs)^26^ (Fig. 1b). The siroheme/Fe_4_S_4_ cluster joins the two S/NiRRs and a long linker mimics the metal-containing co-enzymes in the pseudo-symmetric position on the other face of the SiRHP core^23^. The N-terminal 80 residues of SiRHP are either proteolytically removed or disordered in the crystal structure of the *E. coli* homolog, which is the only known structure of an NADPH-dependent assimilatory SiRHP homolog^23^. In the resting, oxidized state of SiRHP, a phosphate anion binds to the siroheme iron. The nature of the anion was first predicted by enzymology because of a redox-dependent lag in SiRHP activation^27^ and then confirmed with biochemical binding experiments^25,28^. Upon reduction, the phosphate is released through “reduction gated ligand exchange” so that the sulfite substrate binds through its central sulfur^24^.

Here, we show the first near-atomic, high-resolution cryogenic electron microscopy (cryo-EM) structure of the minimal SiRFP/SiRHP dimer, which elucidates their binding interface to understand how SiRHP tightly binds the FNR domain of SiRFP.

## Results

SiRFP largely shares homology with other diflavin reductases like the monomeric CPR in its flexible, three domain architecture with a variable N-terminus. In the case of SiRFP, the N-terminus is solely responsible for oligomerization, followed by an FMN-binding Fld domain, a 36-amino acid long linker, and an FAD/NADPH-binding FNR domain with the intervening connection domain^29^ (Fig. 1). SiRHP is somewhat unique because it shares its siroheme-Fe_4_S_4_ cluster containing active site only with other siroheme-dependent sulfite or nitrite reductases^26^. The core of the monomer has a single active site but shares pseudo-twofold symmetry with heterodimeric dissimilatory homologs, relating two S/NiRRs through a parachute domain that helps form the anion binding cavity^23^ (Fig. 1). In NADPH-dependent assimilatory SiRHPs, the N-terminus is solely responsible for tight binding with its SiRFP partner^3^.

### The SiRHP-SiRFP interaction is highly sensitive to cryo-EM preparation

We determined the structure of the *Escherichia coli* K-12 SiRFP/SiRHP dimer from three modified, minimal dimers, each of which is named by the change to SiRFP and its resulting molecular weight (Figure 1 and Tables 1 and S1). First, we truncated SiRFP to remove the N-terminal Fld domain (SiRFP-43/SiRHP). This is an inactive dimer, because the Fld domain is required for electron transfer. Nevertheless, the two subunits bind tightly and this is the most simplified complex between SiRFP and SiRHP^30^. Second, we truncated both the N-terminal octamerization domain of SiRFP as well as the linker between the Fld and FNR domains to create a monomeric SiRFP that can be locked in an open position (SiRFP-60Δ/SiRHP)^31^. Third, we generated a variant of monomeric SiRFP-60 lacking reactive cysteines into which we engineered a disulfide bond between the Fld and FNR domains but maintained a full-length linker between the SiRFP domains (SiRFP-60X/SiRHP), previously described only in full-length octameric SiRFP^32^. None of the minimal SiR dimers complements SiRFP deficient *E. coli*^33^ (Fig. S1), thus our analysis focuses on the unique, structural interface between the subunits rather than the transient, functional interface that enables electron transfer.

Each variant is highly sensitive to traditional blotting/plunge-freezing methods for cryo-EM preservation. To overcome this sensitivity, we combined the protection of a high critical micelle concentration, fluorinated lipid, fos-choline-8 (FF8, Creative Biolabs, Shirley, NY, USA )^34^, with the blot-free, rapid plunging afforded by the chameleon® system (SPT Labtech, Melbourn, UK)^35^. This cryo-EM sample preparation helped us to retain each intact complex within near ideal ice thickness and avoid denaturation at the air water interface (Fig. S2). The smallest complex (SiRFP-43/SiRHP) showed well-aligned 2D class averages, however the 3D structure revealed structural anisotropy, either due to its small size/asymmetric geometry or from a preferred orientation, that limited high-resolution analysis despite the absence of mobile elements (Figs. S2A and S3). SiRFP-60Δ/SiRHP showed moderate-resolution density (3.54 Å) for the SiRFP FNR domain and SiRHP, however the N-terminal Fld domain was not visible (Figs. S2B and S4A). The 2.78 Å-resolution structure of SiRFP-60X/SiRHP revealed the most detail for the Fld and FNR domains from SiRFP, despite a lack of density for the linker between them in the highest-resolution reconstruction (Figs. 2A, S2C, and S4B). High resolution features for each of the cofactors in both subunits supported this reported resolution (Fig. S5). Therefore, we analyzed the SiRFP-SiRHP interface for this construct in detail.

**Figure 2:**
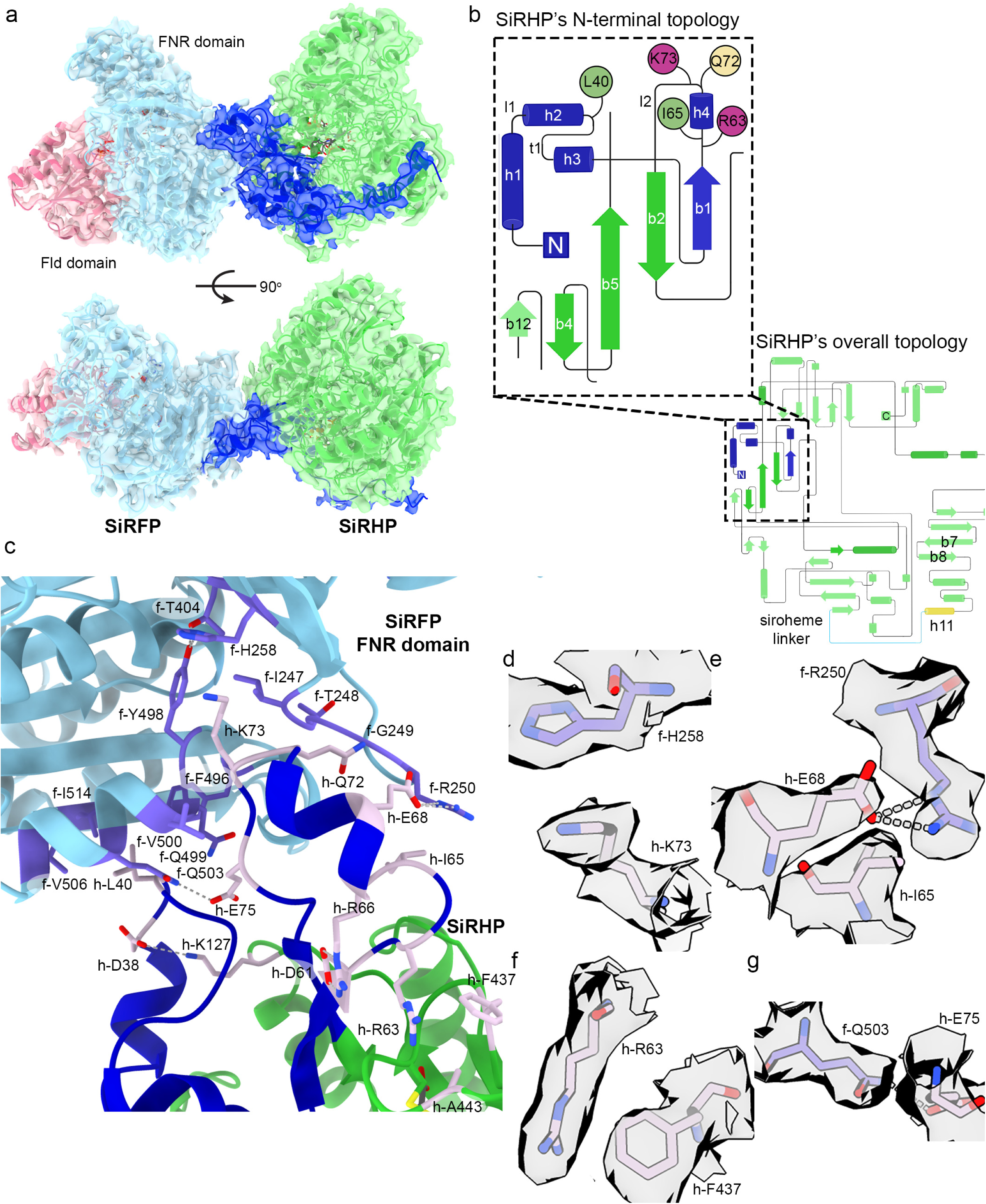
SiRFP and SiRHP interact through the SiRFP FNR domain and the SiRHP N-terminus. **a** The SiRFP/SiRHP interface is minimal, governed by residues from three loops in SiRHP that fit into pockets on SiRFP’s FNR domain. SiRFP’s Fld domain is pink, SiRFP’s FNR domain is light blue. SiRHP’s N-terminal 80 residues are dark blue. SiRHP’s core two S/NiRRs are green. The map is shown at contour level 0.3. **b** The topology diagram of SiRHP, highlighting the N-terminal 80 residues whose structure was previously unknown but that govern the interactions with SiRFP. The residues that interact with SiRFP are localized to three loops. Hydrophobic residues are green, cationic residues are purple, and polar residues are yellow. The first S/NiRR is dark green whereas the second S/NiRR is light green, with the connecting h11 and linker in yellow and cyan, respectively (“Created with BioRender.com.”). **c** The interface between SiRFP and SiRHP is minimal. The SiRFP-FNR domain is light blue with light purple highlighted residues that are important for forming the SiRHP binding site. SiRHP’S N-terminal 80 residues are blue and the S/NiRR repeats are green with mauve highlighted residues that mediate the interaction with SiRFP. **d-g** Select densities for important side chain interactions that mediate the interface between the SiRFP and SiRHP interface are rendered at contour level 0.4 and colored as in **c**.

### SiRFP-SiRHP binding

The SiR dodecameric holoenzyme is composed of four dimers discussed here, when the SiRFP subunit contains the N-terminal octamerization domain, along with four free SiRFP subunits, and is about 800 kDa in mass. Despite this large mass, the binding interface captured in this minimal SiRFP/SiRHP dimer is small, reaching the surface area of 1,138 Å^2^ relative to the overall surface of 43,610 Å^2^. SiRHP alone has a solvent-exposed surface of 25,680 Å^2^. SiRFP-60X alone has a solvent-exposed surface of 21,930 Å^2^. That is, for a large complex only about 2.6% of the solvent-exposed surface is buried upon subunit binding. This is consistent with hydrogen/deuterium exchange-mass spectrometry (H/DX-MS) data on the complex that reveals single, short peptides from each subunit that become occluded upon binding^3^.

The interface is governed by the N-terminus of SiRHP. The structure of this region is previously uncharacterized as it is proteolytically removed in the X-ray crystal structure of *E. coli* SiRHP^23^. These 80 residues follow the topology helix 1 - loop - helix 2 - turn - helix 3 - β-strand 1 - helix 4 - loop - β-strand 2 (Fig. 2b). Only residues from the turn, helix 4, and surrounding loops directly interact with SiRFP. The regions that are N-terminal to the interface interact with domain 1 or the N-terminal half of the parachute domain (*i.e.* the first sulfite or nitrite reductase repeat (S/NiRR)^23,26^), breaking the pseudo two-fold symmetry of SiRHP (Figs. 1b, 2a, and S6a). An extension to the parachute domain within S/NiRR 1 that is not ordered in the original crystal structure, from residues 184-209, helps to hold the N-terminus in place (Fig. S6a). Those residues that are N-terminal to the interface approach the distal active site, but do not contribute significantly to anion or siroheme binding as they are held back by interactions to SiRFP, discussed below. The peptide then turns to form the loop that binds a pocket in SiRFP before moving away from SiRFP in an extended conformation. The N-terminal most residues reach all the way to the other side of SiRHP, interacting with the N-terminus of the α-helix (h11) that precedes the linker that joins the two S/NiRRs and mimics the siroheme binding site (Fig. S6b)^23^.

One interaction that pegs the subunits together is a π-cation interaction between h-Lys73 from SiRHP (for simplicity, residues from SiRHP will be designated with the prefix “h-“) and f-His258 from SiRFP (similarly, residues from SiRFP will be designated with the prefix “f-“) (Figs. 2C and D). This interaction is buttressed by f-Phe496 and f-Val500, which have previously been shown essential for SiRFP-SiRHP binding – the F496D alteration abrogates SiRHP binding and the V500D alteration reduces SiRHP binding 300-fold but both are able to reduce a *cytochrome c* substrate when supplied with NADPH^15^. Further hydrophobic and π-stacking interactions dominate the interface. For example, h-Leu40 inserts into a pocket in SiRFP formed between f-β-sheet 17 and f-α-helix 18, which includes f-Phe496, described above, as well as f-Gln503Cβ, f-Val506, and f-Ile517. h-Ile65 Cγ2 sits 3.3 Å from the plane of the f-Arg250 guanidinium group, which is rotated 90° from its position in free SiRFP^31^ and stabilized by h-Glu68Oε1 (Figs. 2c and e). h-Gln72Cγ also packs into a pocket formed by the backbone atoms of f-Ile247 and f-Thr248, pinned in place by the h-Ile65/h-Glu68/f-Arg250 and h-Lys73/f-His258 interactions. Farther from the interface, there is another stacking interaction between the guanidinium group from h-Arg63 and the h-Phe437 aromatic ring that stabilizes the deformed helix that includes h-Ile65 (Figs. 2c and f). h-Phe437 is adjacent, through h-A443, to the SiRHP iron-sulfur cluster. In this way, the N-terminus of SiRHP, which mediates tight binding to SiRFP, is linked to the siroheme-Fe_4_S_4_ cluster-containing active site through a long-range network of hydrophobic interactions (Fig. S7). The subunits have previously determined to bind tightly, with a *K*_d_ of about 3 nM^15^

Ionic interactions and hydrogen bonds further play indirect roles in the interface by stabilizing the residues and structural elements that mediate the interface (Figs. 2c and S7). For example, a hydrogen bond network from f-Thr404Oγ through f-Tyr498OH and finally to f-His258Nδ1 positions its imidazole ring for the interaction with h-Lys73. An ionic interaction between h-Lys127Nς and h-Asp38Oδ2 reaches across the loop, presenting h-Leu40 to project into a surface pocket on SiRFP. An additional ionic interaction between h-Asp61δ1 and h-Arg66NHδ also stabilizes the deformed helix that contains h-Ile65 and turn it towards f-Arg250.

### The stable interaction between SiRHP and SiRFP differs from the SIR interaction with Fd

In contrast to γ -proteobacteria that couple SiRFP to SiRHP, other organisms like the higher plants *Zea mays* and *Spinacia oleracea* and the actinobacterium *Mycobacteria tuberculosis* use a Fd as their electron carrier partner^19,21^. In those systems, the interaction between the SiR hemoprotein and the Fd is transient whereas there is an additional stable, structural interaction between SiRFP and SiRHP. There is currently no experimentally determined structure of the *M. tuberculosis* SirA bound to its Fd partner. In *Z. mays* SIR, Fd bridges from C-terminal domain 2 to a loop between the first two β-strands (residues Asp110 to Asn118), positioning the Fd iron-sulfur cluster near the SIR metal sites^36^. The interaction is bolstered by *en face* stacking between Fd Tyr37 and SIR Arg324 (Fig. 3a).

**Figure 3:**
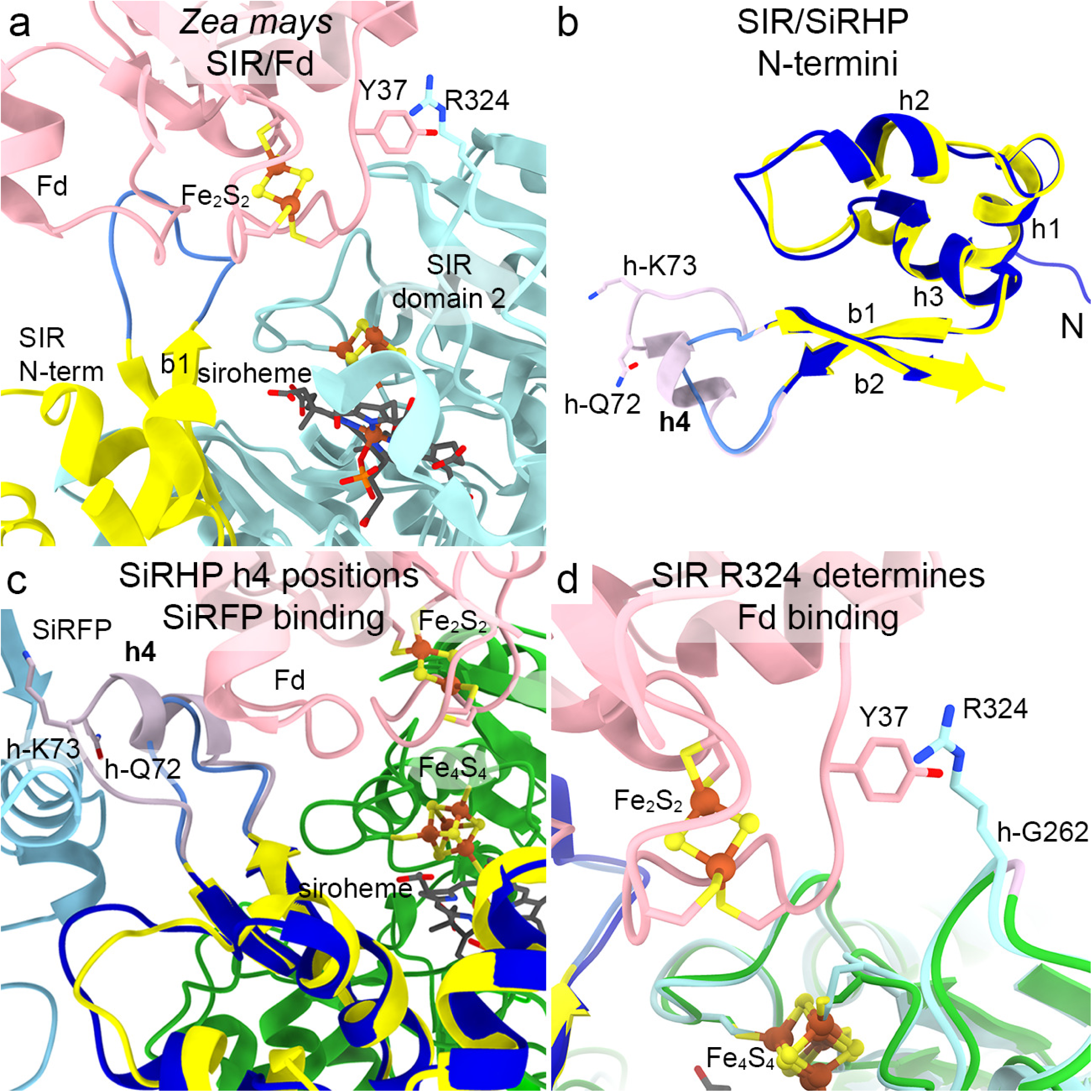
*Zea mays* SIR and *Escherichia coli* SiRHP interact with their reductase partners in different ways. **a** SIR binds a ferredoxin (Fd, pink) such that its Fe_2_S_2_ cluster is close to SIR’s Fe_4_S_4_ cluster, positioned between SIR’s N-terminus (yellow) and domain 2 (cadet blue). (PDB 5H92^36^) **b** SiRHP h4 (mauve) is the sole secondary structural feature that discriminates between SiRFP binding and Fd binding in *Z. mays*. SiRHP’s N-terminal 80 residues are dark blue. SIR’s N-terminal residues are yellow with the element equivalent to SiRHP h4 in medium blue. **c** SiRFP (light blue) and Fd (pink) bind on different faces of SiRHP (dark blue) or SIR (yellow). Only the core S/NiRRs from SiRHP (green) are shown for clarity. Other elements are colored as in **b**. **d** SIR R324 (cadet blue) interacts with Fd Y37 (pink). That interaction is prevented in SiRHP (green) because the position equivalent to SIR R324 is h-G262 (mauve). *Z. mays* Fd Fe_2_S_2_ and SIR Fe_4_S_4_ are shown as in panel **a**.

To understand the structural elements that discriminate between the tight, structural binding interface between SiRHP and SiRFP and the transient interaction between *Z. mays* SIR and Fd, we compared our novel structure with the only known structure of a dimeric assimilatory hemoprotein/Fd structure^20,36^. In SiRHP, the equivalent element between the structurally conserved β-strands 1 and 2 is considerably longer, stretching from h-Asp62 to h-Arg77 and containing a short helix from h-Arg63 to h-Glu71, all found within the least sequence conserved N-terminus (Figs. 3b and S8). This loop contains h-Gln72 and h-Lys73 that help form the interaction with FNR domain in SiRFP, which faces away from where Fd binds to the *Z. mays* homolog (Fig. 3c). Further, the arginine in SIR that stacks with Fd Tyr37 is not conserved in SiRHP – the equivalent position is h-Gly262, despite the structural conservation of the loop between β-strands 7 and 8 that contributes to siroheme binding (Fig. 3d).

Interestingly, the Fd-dependent *M. tuberculosis* SirA structure, which was initially mis-identified as a siroheme-dependent nitrite reductase^20^, has a lysine at the equivalent position to h-Gly262 but shares the longer insertion with SiRHP (Fig. S8a). In that structure, which does not have a bound Fd, however, the helical element points away from either SiRFP, in the *E. coli* structure, and sterically clashes with the Fd as bound in the *Z. mays* structure (Fig. S8b). In this way, that divergent structural loop plays a role in electron transfer partner binding, whether stably, as in the interaction between SiRHP and the SiRFP FNR domain; transiently, as in *Z mays* SIR/Fd; or to determine Fd specificity in an uncharacterized way, as in *M. tuberculosis* SirA/Fd.

### The SiRHP active site loop is in its anion-binding conformation

When bound to SiRFP, the anion binding loop in SiRHP (h-Asn149 to h-Arg153) is in its closed position, held in place by a long (3.9 Å), through-space interaction between h-Arg53Cδ and h-Asn149Cβ (Fig. 4a). h-Arg53 is, in turn, held in place by stacking between its guanidinium group and h-Tyr58. The ring of h-Tyr58 subsequently sits over the methyl group on the siroheme pyrroline A ring. The only other new protein/siroheme interaction identified in this now complete structure of SiRHP is an ionic bond between h-Gln60 and the propionyl group from siroheme pyrroline ring B. The siroheme is saddle-shaped, as in free SiRHP and unlike in dissimilatory SIR^37–40^. h-Arg153 is flipped away from the bound phosphate. The other three anion binding residues, h-Arg83, h-Lys215, and h-Lys217, remain largely unchanged from free SiRHP (Figs. 4a-c)^23^.

**Figure 4:**
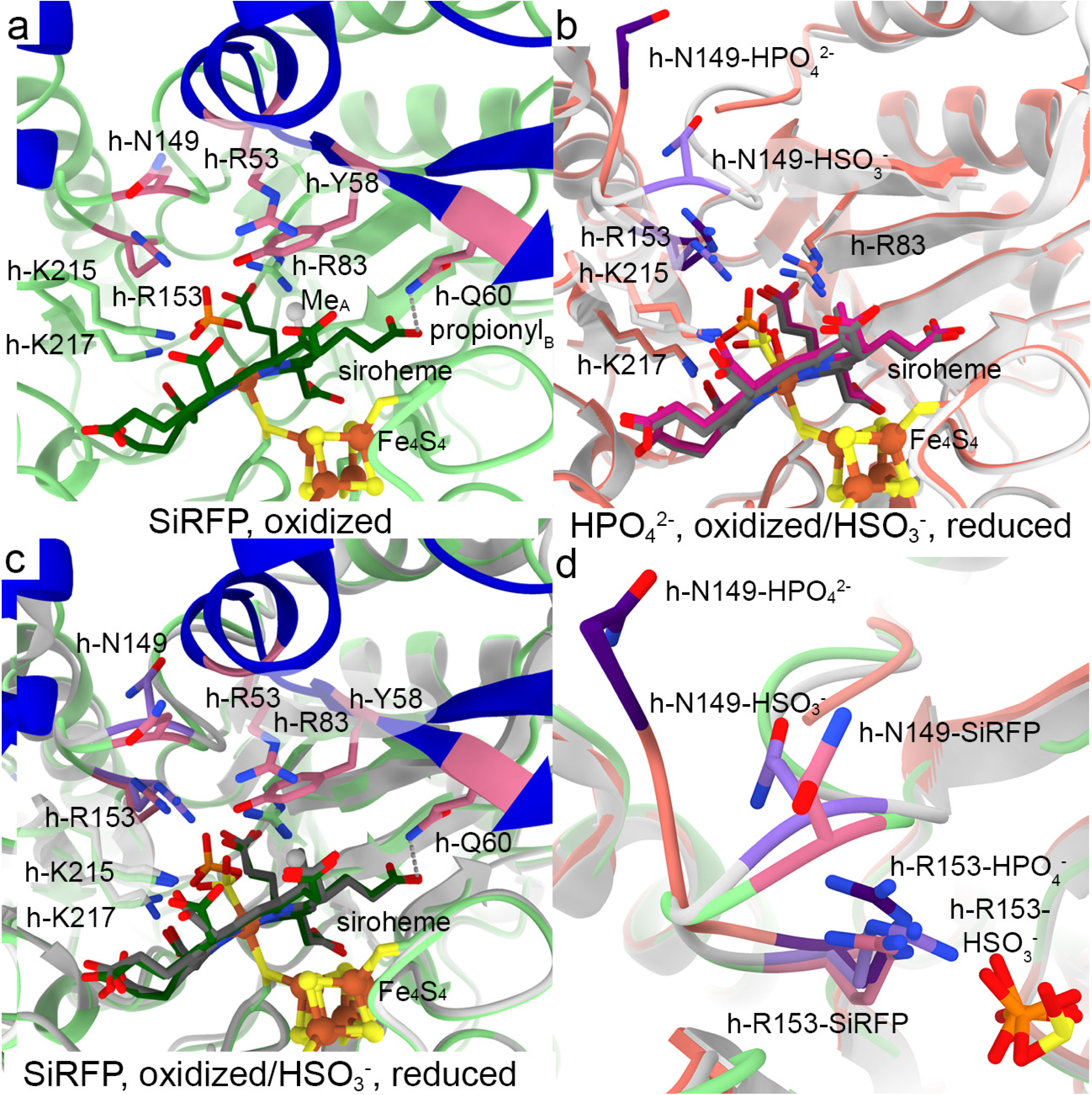
SiRHP’s active site loop that forms the complete substrate binding site is in its closed conformation with the intact N-terminus, bound to SiRFP. **a** When SiRFP binds oxidized SiRHP with the bound, inhibitory phosphate, the anion-binding loop containing h-N149 is ordered but with h-Arg153 in a similar orientation as in free SiRHP. Panels are labeled by their binding partner and oxidation state. Residues important for anion binding or loop position are dark pink. The siroheme methyl group from ring A is a gray sphere and the interaction between h-Q60 and the ring B propionyl group is marked by a gray dash. **b** In the absence of its N-terminal 80 residues, SiRFP, or the sulfite substrate, SiRHP’s anion binding loop is disordered (missing salmon ribbons marked by salmon *s). The residues that are important for anion binding in response to oxidation state in this structure are dark purple, those that are unchanged are salmon (PDB 1AOP^23^). Upon reduction and with bound sulfite, the active site loop becomes ordered and h-R153 flips over to bind the substrate (light gray ribbons). The residues that are important for anion binding in response to oxidation state in this structure are light purple, those that are unchanged are light gray (PDB 2GEP^24,25^). **c** Superimposition of the SiRFP-bound SiRHP (green) and the reduced, sulfite-bound, SiRHP (light gray) shows the anion-binding loop to be in similar position as in reduced/SO_3_^2-^-bound SiRHP, primed for sulfite binding. **d** Superimposition of the three SiRHP structures bound to different anions or binding partner highlights the intermediate conformation induced by SiRFP binding. This intermediate conformation shows ordering of SiRHP’s N-terminus and the active site loop in a phosphate-bound, oxidized state. Models are colored as in **a-c**.

This conformation differs from the various redox and anion-bound structures of free SiRHP^23–25,28^. In the phosphate-bound, free SiRHP structure that lacks the N-terminal 80 amino acid extension^23^, the loop is flipped open such that h-Ala146 through h-Ala148 are disordered (Fig. 4b). Upon reduction and sulfite binding, h-Arg153 flips over to interact with the smaller anion and the loop becomes ordered^25^. h-Asn149 points away from the active site. In this way, SiRFP binding to SiRHP, with the ordering of the SiRHP N-terminus, induces an intermediate structure with elements of both the oxidized, phosphate bound and the reduced, sulfite bound conformations (Fig. 4d).

In the original SiRHP X-ray crystal structure, the siroheme iron is significantly domed above the siroheme nitrogens, indicative of an oxidized Fe^3+^ (Fig. 4b)^23^. Subsequent chemical reduction experiments show the doming flattens upon conversion to Fe^2+^, commensurate with release of the phosphate to allow substrate binding (Fig. 4b)^25^. Contemporary X-ray diffraction experiments show that this reduction is beam-induced, but within the constraints of the crystal the phosphate remains bound to the siroheme iron in the active site^28^. In this cryo-EM structure, the siroheme iron appears to be in the plane of the siroheme nitrogens, suggesting that it has also been reduced by the electron beam. The central density for the phosphate (B-factor 36 Å^2^) is 3.5 Å from the siroheme iron and its pyramidal shape is rotated such that the oxygen-iron bond appears broken (Figs. 4a and S5c).

### SiRFP is highly mobile

Although the 2D class averages in all three datasets appeared to show little orientation preference with discernable features, further analysis revealed each to have unique properties related to the SiRFP variant used to generate the dimer (Table S1, Figs. S2-4).

SiRFP-43/SiRHP (107 kDa in mass): The 2D class averages for SiRFP-43/SiRHP appeared to show high-resolution features, however the initial 3D models were poorly aligned, likely due to a combination of small mass, preferred orientation, and limitations in grid preparation, so the refined volume did not achieve high resolution despite its simplified form (Fig. S3). The absence of the SiRFP Fld domain did not alter the SiRFP-SiRHP interaction, as described below, or the binding of the FAD cofactor as compared to the crystal structures^31,41^.

SiRFP-60Δ/SiRHP (123 kDa): As with SiRFP-43/SiRHP, this dimer showed 2D class averages with high-resolution features for the of the dimer (Figs. S2b and S4a). Nevertheless, the initial models were of inconsistent structure, so we checked for heterogeneity using “3D variability” in CryoSPARC, which revealed a distinctive degree of movement in the Fld domain. To better understand the degree of flexibility for the Fld domain from the SiRFP-60Δ/SiRHP dimer, we performed both “3D flex” in CryoSPARC as well as “heterogenous refinement” in cryoDRGN^42^ with the particles from refinement on the main heterodimer body. This analysis, anchored on the SiRFP-FNR/SiRHP dimer, identified a dramatic movement of the SiRFP Fld domain relative to the FNR domain, swinging 20° between the most compact and most open forms (Video S1 and Figs. 5A and S9). The density for the linker joining the domains is not visible. In its most compact conformation, the Fld domain reaches the canonical “closed” conformation in which the Fld domain tucks into a cavity in the FNR domain, bringing together the FAD and FMN cofactors for electron transfer as in the homologous CPR structure after the FAD is reduced by NADPH (Fig. 5b and S9). In the most open conformation, the Fld domain assumes a different position to that seen in the “open” conformation determined by X-ray crystallography of the same monomeric variant and the average solution envelope determined from small angle neutron scattering (SANS), intermediate between the fully opened and closed conformations (Fig. 5c and S9)^30,31^.

**Figure 5:**
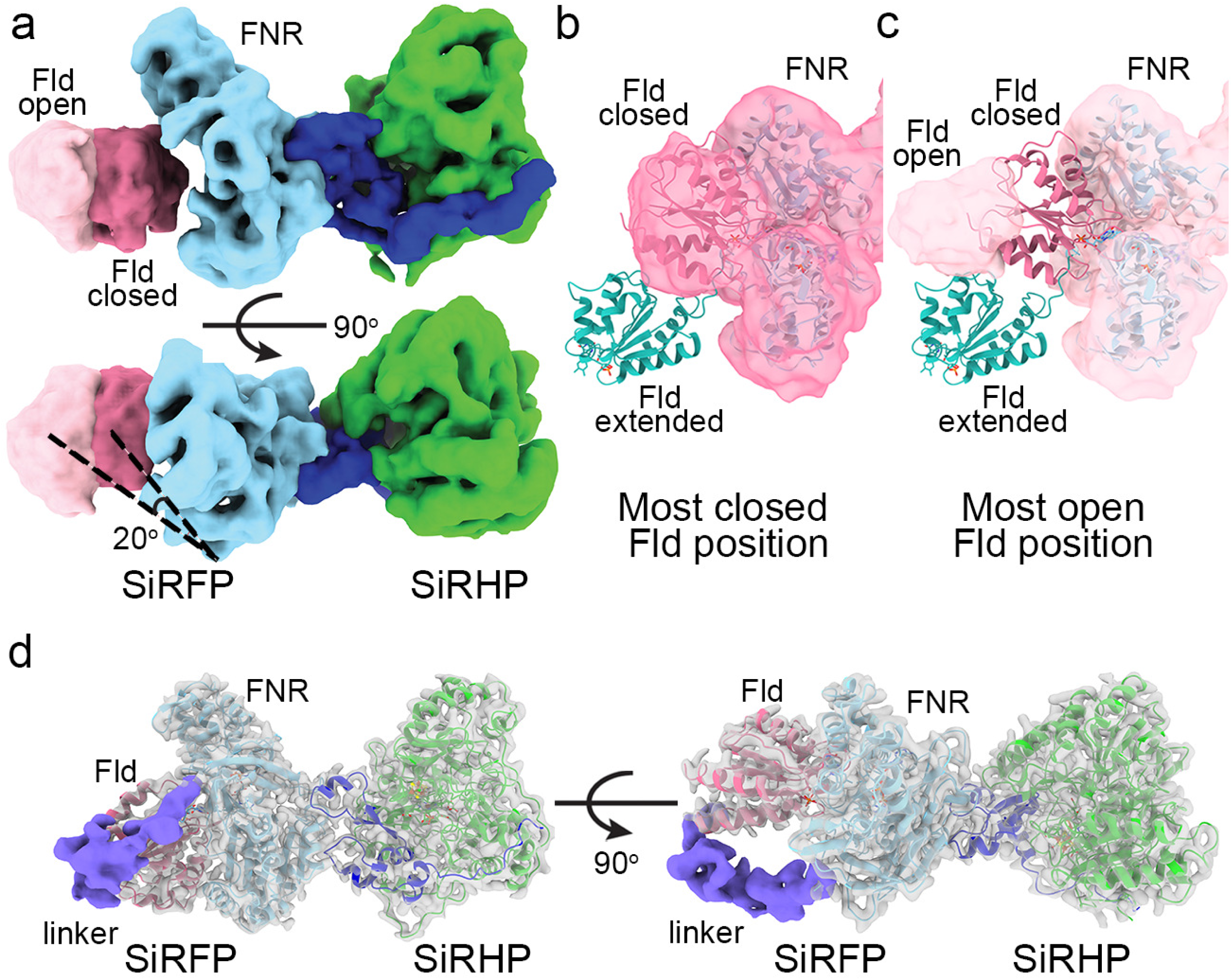
The SiRFP Fld domain is highly mobile relative to its FNR domain. **a** One of the latent variable dimensions of motion demonstrates an opening of the Fld domain, even in SiRFP-60Δ, with the motion parallel to the midline of the FNR domain (light blue). The most-closed position is dark pink, the most open is light pink. SiRHP is green, with the N-terminus in dark blue. The angle is measured between the lines connecting the first traceable residues in the FNR domain with the open and closed Fld volume center of mass. **b** The map of the most closed conformation of the Fld domain (dark pink) corresponds to the position of the Fld domain in the dimer formed with SiRFP-60X, far from where it is in the extended conformation of SiRFP-60Δ in crystals (teal, PDB 6EFV^31^). **c** The map of the most opened conformation (light pink) shows that the Fld domain lands in an intermediary position between a highly extended position seen in the X-ray crystal structure (teal, PDB 6EFV^31^) and the closed conformation seen in **b**. **d** Even with the Fld and FNR domains constrained by a crosslink, the 36 residue-long linker between the domains is largely disordered, visible only in a small (∼20,000) subset of particles (purple volume), shown at contour level 0.06.

SiRFP-60X/SiRHP (124 kDa): The Fld and FNR domains are covalently linked in this dimer, thus restricting the swinging domain motion of the Fld domain seen in the SiRFP-60Δ/SiRHP dimer. No preferred orientation was identified either in 2D or 3D analysis (Fig. S4b). Nevertheless, there is no density for the linker between the Fld and FNR domains in the highest-resolution reconstruction, which suggests that the 36 amino acid linker is exceptionally dynamic but allows the two SiRFP domains to come into close contact to catch them in a crosslink. The Fld domain remains the region with the highest B-factors (Fig. S10). Locking the Fld and FNR domains in the closed conformation resulted in the most stable variant for high-resolution analysis (2.78 Å), discussed above. In SiRFP-60X, the FAD and FMN cofactors bind as they do in the open conformation in X-ray crystal structure of SiRFP-60Δ^31^ (Fig. S11). When compared to the closed conformation formed by crystal contacts in the X-ray crystal structure ^31^, however, the Fld domain loop containing f-Tyr158 shifts about 2.8 Å towards the FNR domain, moving the FMN cofactor about 0.7 Å closer to the FAD (Fig. S11). The C-terminal most f-Tyr599 that stacks with the FMN and is unique to SiRFP compared to CPR^41^, does not change regardless of whether the closed conformation is formed from the crystal contacts or the engineered crosslink (Fig. S11). The relative orientation of the flavins is also not disturbed by the engineered crosslink (Fig. S11), unlike the effect of an analogous crosslink in CPR^43^.

With further classification of the SiRFP-60X/SiRHP structure, a sub-class of ∼20,000 particles (10% of the particles used for highest resolution reconstruction for this dataset) emerged that revealed the linker between the two domains. The linker runs from the C-terminal end of the Fld domain (amino acid f-Ser207) to the N-terminal end of the FNR domain (amino acid f-Ile226) (Fig. 5d). There is neither defined secondary structure nor contact with either of the SiRFP domains, thus the linker is not constrained in a single conformation across the entire ensemble of particles even when the domains are crosslinked. The highly mobile nature of these 36 residues explains how the Fld domain can reposition for contact with the active site containing partner, SiRHP, presumably to mediate electron transfer.

## Discussion

The interface between SiRFP and SiRHP in a minimal heterodimer is surprisingly small, driven by π-cation, hydrophobic, planar stacking between aromatic side chains, and ionic interactions. The non-canonical planar stacking interactions with strategic arginine residues are not as uncommon as would be expected from the canonical role that arginine plays in electrostatic interactions and are often found at important structural elements of the protein or enzyme^44^. In the case of the SiRFP/SiRHP dimer, these interactions either mediate the dimer interface itself or establish the structural platform to position other side chains to mediate the interface. Polar or electrostatic interactions also play supporting roles to align the non-polar interactions because the arginine amino groups are free to make independent, more canonical, interactions with other side chains.

The curious predominance of non-ionic interactions that holds the complex together explains why cryo-EM analysis has been elusive – the hydrophobic air-water interface in a thin film for cryogenic plunging is destructive for such interactions^45–48^. We performed extensive trial-and-error probing grid type, grid substrate, hole size, method for hydrophilizing the grid, detergents, and plunging methods to identify what seems to be a singular condition that allowed cryogenic preservation for cryo-EM analysis: the combination of FF8 and the chameleon® blot-free, rapid plunging. Neither FF8 alone with traditional plunging nor chameleon® alone without FF8 was sufficient. Such low-throughput screening to find the one combination of variables that allows for high-quality sample preparation highlights the need for further technology development on the front-end of cryo-EM structure determination. Simply put, high-throughput data collection with direct electron detectors is not always sufficient to overcome cryo-EM sample preparation challenges.

Surprisingly minimal structural elements determine the evolutionary pressure for siroheme-dependent SiR hemoproteins to bind tightly with a diflavin reductase like SiRFP or transiently to a simpler Fd. For example, in comparing the only known structures of monomeric assimilatory SiR hemoproteins with their electron transfer partners – *E. coli* SiRHP (this work) and *Z. mays* SIR^36^ – there is a loop extension in *E. coli* SiRHP between a conserved two-strand β-sheet that binds to SiRFP, compared to how that loop behaves in the case of *Z. mays*. Additional structural analysis of *M. tuberculosis* SirA or *S. oleracea* SiRHP bound to their unidentified Fd remain to determine how each unique hemoprotein behaves. Of course, there is an analogous interaction between SiRHP and the Fld domain from SiRFP – there must be for electron transfer to occur, but its structure remains elusive because that interaction is so transient in the SiRFP/SiRHP heterodimer that it cannot be captured biochemically^3,15^. Further, how that interaction forms in the context of dodecameric SiR would be challenging to model because the Fld domain is at the core of the SiRFP complex whereas SiRHP is peripheral^32^.

The minimal SiRFP/SiRHP dimeric structure reported here shows that when SiRHP binds SiRFP, the anion-binding active site loop is constrained from disorder, unlike in free SiRHP lacking its N-terminus^23^. Nevertheless, h-Arg153 is in its phosphate-binding conformation. The position of the loop is, thus, impacted by the presence of the SiRHP N-terminus that is absent in the X-ray crystal structure. The importance of this observation is understood in comparison to a series of X-ray crystal structures in various redox states and bound to various anions^24,25^. This series of structures shows how the active site re-orients with the changing state, namely that the loop between h-Asn149 to h-Arg153 is disordered in the oxidized state in the absence of its N-terminus or bound SiRFP partner. Upon reduction the loop becomes ordered and h-Arg153 flips to bind the substrate.

The question that remains is how the SiRFP Fld domain approaches the SiRHP metallic active site for productive electron transfer because that transient interaction does not mediate the structural interface between the subunits^3^. Clearly there is dramatic flexibility within SiRFP – even when the Fld and FNR domains are covalently attached by an engineered crosslink, we cannot visualize the linker between the domains without extensive classification and the Fld domain has high B-factors (Figs. 5D and S10). In the SiR-X dodecameric holoenzyme, we have observed both intermolecular and intramolecular crosslinks^32^. Crystallographic studies of SiRFP-60 show that the position of the Fld domain relative to the FNR domain is not fixed^41^ without further truncation of the linker joining the domains^31^. In solution, as studied by SANS, the relative position of the Fld and FNR domains depends on the oxidation state of SiRFP and whether or not it is bound to SiRHP^30^.

Taking together all of the different positions in which the Fld domain has been identified, the Fld domain can move at least 58 Å from its position in reduced SiRFP-60, modeled from a low-resolution SANS envelope^30^, to its crosslinked position determined from the high-resolution cryo-EM structure presented here, measured from the phosphate of the FMN co-enzyme (Fig. S9a). Using those same phosphates and anchoring at the siroheme iron, the domains move through an angle of 67°, facilitated by the 36-amino acid long linker that joins the Fld and FNR domains (Fig. S9a). Of note, contrast matching SANS experiments on the minimal dimer^30^ supported the prediction by H/DX-MS and mutational analysis^3,15^, placing SiRHP adjacent to the FNR domain. Y101 from the SiRFP Fld domain was identified through H/DX-MS analysis as subject to a change in solvent accessibility upon SiRHP binding, but was not essential for either tight binding or enzyme function^3,15^ Further, SANS experiments on the reconstituted holoenzyme dodecamer revealed that the four SiRHP subunits bind the SiRFP octamer independently and far from the Fld domain^2,32^. None of these low-resolution techniques determined the atomic-resolution interactions that mediate SiRFP-SiRHP binding, described here.

Further comparison of the varying Fld positions with monomeric SiRFP-60 when crystallized^31,41^ highlights the nature of its plasticity that likely underlies its ability to form a transient, functional interface with SiRHP. That is, four of the five crystal structures reported for SiRFP-60 show large solvent channels with no electron density that can be accounted for by the Fld domain; two of those structures are reported in the PDB^41^. In one crystal form, the channels can largely accommodate the Fld domain positions determined from the cryo-EM structure and SANS^30^ models (Fig. S9b). In the other, there is steric clashing with some positions (Fig. S9c). In both crystal forms, the SiRHP binding interface interacts with another SiRFP-60 molecule, suggesting that interface has a propensity to bind other proteins. The model that emerges, which sets SiRFP apart from its CPR homolog, is that the repositioning of the Fld domain to interact with the hemoprotein partner is not a simple opening relative to the FNR domain. Rather, the Fld domain seems to be able to rotate through an elliptical cone-shaped range of positions (Fig. S9d), giving it the flexibility to interact with a tightly bound SiRHP (provided the linker is sufficiently long) or, in the dodecamer, a SiRHP bound to a partner.

Despite our extensive efforts at determining the high-resolution structure of the full complex, the dodecamer has proven to be highly heterogeneous, to date eluding the power of contemporary heterogeneity analysis of cryo-EM images. The basis for the extreme flexibility of SiRFP is the 36 amino acid long linker between the Fld and FNR domains, compared to a 12 amino acid long linker in CPR^43^. Heterogeneity analysis of SiRFP-60Δ in complex with SiRHP identified a dominant vector of motion parallel to the major axis of the FNR domain. This is in contrast to the highly extended conformation seen in the X-ray crystal structure of free SiRFP-60Δ^31^ and the various positions of the Fld domain determined by resolution SANS envelope models of SiRFP-60 in various states (Fig. S9). These different experimentally determined structures do not identify a single axis along which the Fld domain moves relative to the FNR domain to show how the Fld domain makes a transient interaction with SiRHP for electron transfer. Nevertheless, prior biochemical experimentation shows that cross-subunit interactions occur within the SiRFP octamer^32^. These observations support the hypothesis that within the holoenzyme, there is not a singular inter-subunit interaction between the Fld and FNR domains of any given SiRFP subunit within the octamer to allow reduction of the FMN cofactor. Thus, a distinct, transient, and electron transfer mediating interaction between a single SiRFP Fld domain and SiRHP partner does not likely exist within the full, dodecameric holoenzyme complex.

## Conclusions

The combined technological developments in cryo-EM over the past decade all played a role in determining the structure of this elusive complex, from cryogenic sample preservation to data collection to image analysis. In doing so, we show that the subunits of NADPH-dependent assimilatory sulfite reductase bind through an interface governed by the N-terminus of SiRHP and the FNR domain of SiRFP. The interaction is surprisingly minimal, governed by a set of hydrophobic and ionic interactions. Structural pairing between a siroheme-dependent hemoproteins and diflavin reductase or Fd appears to fall to a short loop between a conserved 2-stranded β sheet. The high mobility of the SiRFP Fld domain relative to its FNR domain likely explains how a minimal dimer maintains redox-dependent functionality with a full-length linker. Understanding the flexibility that stems from the linker between the SiRFP Fld and FMN domains, even when the linker is shortened or the domains are crosslinked, suggests a new bottleneck to be overcome in determining the structure of the full SiR holoenzyme.

## Materials and methods

### Protein Expression and Purification

Each *E. coli* K12 SiRFP/SiRHP dimer variant was generated and purified as previously described^2,3,15,30,32^. The SiRFP or SiRHP-expressing pBAD constructs^3,15,30^ were subcloned from pJYW632^49^ *cysJ* or *cysI*/*cysG* (SiRFP: UniProtKB accession code P38038 and SiRHP: UniProtKB accession code P17846). Briefly, untagged SiRHP was co-purified with the following SiRFP variants: 1) SiRFP-43, a 43 kDa monomeric SiRFP variant lacking the N-terminal Fld domain and linker^30^; 2) SiRFP-60Δ, a 60 kDa monomeric variant of SiRFP generated by truncating its first 51 residues with an additional internal truncation of six residues (Δ-AAPSQS) in the linker that joins the Fld and FNR domains^31^; and 3) SiRFP-60X, the 60 kDa monomer with an engineered disulfide crosslink between the Fld and FNR domains, as has previously been analyzed in octameric SiRFP^32^. In SiRFP-60X, two background cysteines (C162T and C552S) were altered to avoid unwanted crosslinking and two cysteines were added (E121C and N556C) to form a disulfide bond between Fld and FNR domains of SiRFP but with a full-length linker between the domains. DNA sequencing confirmed the presence of all deletions and mutations in the variants. All related SiR constructs are available from the corresponding author upon request.

*E. coli* LMG194 cells (Invitrogen, Carlsbad, CA, USA) were transformed with the pBAD plasmid containing the genes encoding SiRFP-60Δ, SiRFP-60X, SiRFP-43 or SiRHP. SiRHP and SiRFP variants were expressed independently. The SiRFP-43/SiRHP and SiRFP-60Δ were assembled by mixing cells prior to lysis, and the formed heterodimers were co-purified. The SiRFP-60X/SiRHP dimer was assembled by reconstituting SiRFP-60X with 2 Eq SiRHP for 1 h on ice and isolated as before for similar dimers^2,30,32^. An N-terminal six-histidine tag was present in all SiRFP variants whereas SiRHP was untagged. Each recombinant *E. coli* strain was grown and induced at 25 °C with 0.05% L-arabinose. SiRFP-43, SiRFP-60Δ and SiRHP cells were expressed for 4 hr whereas SiRFP-60X was expressed overnight. All variants were purified using a combination of Ni-NTA affinity chromatography (Cytiva, Marlborough, MA, USA), anion exchange HiTrap-Q HP chromatography (Cytiva, Marlborough, MA, USA) and Sephacryl S300-HR size exclusion chromatography (Cytiva, Marlborough, MA, USA) using previously optimized SPG buffers (17 mM succinic acid, 574 mM sodium dihydrogen phosphate, pH 6.8, 374 mM glycine, 200 mM NaCl)^2,15,30,32^. All variants have been extensively characterized with UV-Vis spectroscopy, size exclusion chromatography, SDS PAGE analysis for correct stoichiometry (Fig. S12) and with SANS for solution monodispersity^2,15,30,32^. This is the first analysis of the heterodimeric SiRFP-60X/SiRHP assembly.

### SiRFP-60 variant complementation analysis

SiRFP-deficient *E. coli* (*cysJ*^-^, Keio strain JW2734^33^) cells were separately transformed with plasmids expressing SiRFP-60, empty pBAD vector, SiRFP-60Δ, SiRFP-60X, or SiRFP-43. Cells were grown overnight in Luria-Bertani (LB) medium with 50 μg/mL kanamycin and 100 μg/mL ampicillin. All cultures were harvested and washed gently three times in M9 salts. Cell densities were normalized at 0.45 OD_600_ before plating serial dilutions onto either M9-agar plates containing 50 μg/mL kanamycin, 100 μg/mL ampicillin, or LB media with the same antibiotics. Kanamycin and ampicillin maintained the *cysJ*^-^ deficiency and the pBAD plasmid, respectively. Cell growth was assessed after 48 hours.

### Cryo-EM sample preparation

SiRFP-43/SiRHP, SiRFP-60Δ/SiRHP and SiRFP-60X/SiRHP were each plunged using the chameleon® blotless plunging system (SPT Labtech, Melbourn, UK) at 10 mg/mL protein. chameleon® self-wicking grids^35^ were backed with 18-gauge gold (Ted Pella, Redding CA, USA) on an Auto 306 vacuum coater (BOC Edwards, West Sussex, UK). The following glow discharge (GD)/wicking times (WTs) were used: SiRFP-43/SiRHP: 30 s GD, 130 ms WT; SiRFP-60Δ/SiRHP: 80 s GD, 195 ms WTs; SiRFP-60X/SiRHP 45 s GD, 175 ms WTs. All samples were premixed with fluorinated FC-8 detergent^34^ at 2 mM final concentration. The plunged grids were clipped and then screened for high quality with a Titan Krios operating the Leginon software package^50^.

### Data collection and processing

1. SiRFP-43/SiRHP: 14,600 movies were collected at 300 KV on a Titan Krios using a GATAN K3 camera with 0.844 Å/pixel and the Leginon automated data collection package^51,52^. After motion correction with Motioncor2^53^ in the Relion GUI^54^, CTF estimation was performed using CTFFIND4^55^. Particles were picked by “blob picker” followed by “template picker” in CryoSPARC^56^. Initial 2D classification, followed by multiple rounds of 2D class selection/classification, identified 1,500,000 particles that were used for initial map building and non-uniform refinement in CryoSPARC. This process achieved a reported 3.6 Å-resolution map of the minimal dimer. However, the map features did not reflect the reported resolution. 3D variability analysis in CryoSPARC^56^ did not reveal significant conformational mobility, however orientation analysis and calculation of the 3DFSC^57^ showed the particles harbored a preferred orientation (Fig. S3). “Orientation diagnostic” in CryoSPARC was performed to confirm this interpretation. To further confirm and diminish the preferred orientation issue, “Rebalance 2D” was performed to put particles into 7 super classes and limit the total particles in each superclass down to total of 350,000 and 105,000 particles. Non-uniform refinement^58^ followed by orientation diagnostics performed in CryoSPARC produced a more accurate resolution (4.31 Å, 4.74 Å respectively) and better quality map. deepEMhancer^59^ was used to sharpen these maps (Fig. S2). Particle picking performed by the TOPAZ algorithm^60^ gave the same result.
2. SiRFP-60Δ/SiRHP: 25,488 movies were collected at 300 KV on a Titan Krios using a GATAN K3 camera with 0.844 Å/pixel and the Leginon automated data collection package^51,52^. Motion correction, CTF estimation, and particle picking were performed as for SiRFP-43/SiRHP. After multiple rounds of 2D classification, ∼550,000 particles were used for initial map building and non-uniform refinement in CryoSPARC to achieve the final 3.49 Å-resolution structure of the dimer, masked to omit the SiRFP Fld domain. Multiple refinements with different masking were performed. Masking to include the whole complex, including the SiRFP Fld domain, resulted in a low-resolution reconstruction for the Fld domain. Particles were then down sampled and imported to cryoDRGN to perform heterogenous refinement, giving a series of volumes showing the Fld domain movement. CryoSPARC 3D Flex^61^ gave the same result as cryoDRGN.
3. SiRFP-60X/SiRHP: ∼10,000 movies were collected at 300 KV on a Titan Krios using a DE Apollo camera with 0.765 Å/pixel and the Leginon automated data collection package^51,52^. Motion correction, CTF estimation, and particle picking were performed as above. Multiple rounds of 2D classifications identified ∼185,000 particles that were used for initial map building and non-uniform refinement in CryoSPARC to achieve a 2.84 Å-resolution structure of the entire dimer, including the SiRFP Fld domain. Due to the GFSC curve not going to zero, bigger box size was used for the particle extraction (360 pixels, previously 320) and ran into the job “Remove Duplicates” in CryoSPARC to remove the particles closer than 150 Å to each other. These two steps reduced the number of particles to 179,100 and the resolution improved to 2.78 Å with a GFSC curve becoming closer to zero. Further 3D variability was performed to track the Fld linker. After multiple rounds of 3D classification, 10% of the particles (∼20,000) were used to perform a non-uniform refinement, resulted in a 3.61 Å overall resolution structure, sharpened by deepEMhancer including the linker.

### Model building and refinement

Model building was initiated using the “fit in map” algorithm in Chimera^62^ using the atomic model of SiRFP from PDB ID 6EFV^31^, fitting each of the Fld or FNR domains independently or the atomic model of SiRHP generated in AlphaFold^63,64^ to capture its N-terminal 80 residues. Iterative real-space refinement in PHENIX^65^ with manual fitting in Coot^66^ yielded a model with a correlation of 0.85 (Table S3). The model for SiRFP-60X/SiRHP was deposited in the PDB (https://www.rcsb.org) as 9C91 and the map for SiRFP-60X/SiRHP and the related map series for SiRFP-60D/SiRHP was deposited in the EMDB (https://www.ebi.ac.uk/emdb/) as EMD-45359. The data in the form of the particle image stack used for the final 3D reconstruction was deposited in the EMPIAR database under the accession number EMPIAR-12180 (https://www.ebi.ac.uk/empiar/).

## Acknowledgements and funding

We kindly thank Dr.s Scott Stagg, Qian Yin, and Christopher Stroupe for helpful discussions. Some of this work was performed at the Simons Electron Microscopy Center at the New York Structural Biology Center, with major support from the Simons Foundation (SF349247). Florida State University supports cryo-EM in the Biological Imaging Resource Center, which houses the following equipment used in this study: a Gatan Solaris Plasma Cleaner (NIH grant S10 RR024564), a Hitachi HT7800 (NSF grant MRI2017869), a ThermoFisher Vitrobot Mark IV (NIH grant S10 RR024564), an SPI chameleon® plunging system (NIH grant R24 GM145964), a ThermoFisher Titan Krios (NIH grant S10 RR025080), and a DE Apollo direct electron detector (NIH grant R35 GM139616). This work was further supported by National Science Foundation grants MCB1856502 and CHE1904612 to M.E.S.

## Author contributions

BGE participated in data collection and performed the data analysis. NW and BGE prepared the protein specimens. YG performed the complementation analysis. KN and MA prepared cryogenic grids and participated in data collection. IA optimized protein specimen preparation. HAW and JHM supervised cryogenic specimen preparation and data collection. MES conceived of the study and oversaw experimentation.

## References

1 Siegel, L. M. & Davis, P. S. Reduced nicotinamide adenine dinucleotide phosphate-sulfite reductase of enterobacteria. IV. The *Escherichia coli* hemoflavoprotein: subunit structure and dissociation into hemoprotein and flavoprotein components. J Biol Chem 249, 1587–1598 (1974).

2 Murray, D. T. et al. Neutron scattering maps the higher-order assembly of NADPH-dependent assimilatory sulfite reductase. Biophys J 121, 1799–1812 (2022). 10.1016/j.bpj.2022.04.021

3 Askenasy, I. et al. The N-terminal Domain of *Escherichia coli* Assimilatory NADPH-Sulfite Reductase Hemoprotein Is an Oligomerization Domain That Mediates Holoenzyme Assembly. J Biol Chem 290, 19319–19333 (2015). 10.1074/jbc.M115.662379

4 Siegel, L. M., Davis, P. S. & Kamin, H. Reduced nicotinamide adenine dinucleotide phosphate-sulfite reductase of enterobacteria. 3. The *Escherichia coli* hemoflavoprotein: catalytic parameters and the sequence of electron flow. J Biol Chem 249, 1572–1586 (1974).

5 Wang, M. et al. Three-dimensional structure of NADPH-cytochrome P_450_ reductase: prototype for FMN- and FAD-containing enzymes. Proc Natl Acad Sci U S A 94, 8411–8416 (1997).

6 Girvan, H. M. et al. Flavocytochrome P_450_ BM3 and the origin of CYP102 fusion species. Biochem Soc Trans 34, 1173–1177 (2006). 10.1042/BST0341173

7 Garcin, E. D. et al. Structural basis for isozyme-specific regulation of electron transfer in nitric-oxide synthase. J Biol Chem 279, 37918–37927 (2004). 10.1074/jbc.M406204200

8 Zhang, J. et al. Crystal structure of the FAD/NADPH-binding domain of rat neuronal nitric-oxide synthase. Comparisons with NADPH-cytochrome P_450_ oxidoreductase. J Biol Chem 276, 37506–37513 (2001). 10.1074/jbc.M105503200

9 Olteanu, H. & Banerjee, R. Human methionine synthase reductase, a soluble P-_450_ reductase-like dual flavoprotein, is sufficient for NADPH-dependent methionine synthase activation. J Biol Chem 276, 35558–35563 (2001). 10.1074/jbc.M103707200

10 Freeman, S. L., Martel, A., Raven, E. L. & Roberts, G. C. K. Orchestrated Domain Movement in Catalysis by Cytochrome P_450_ Reductase. Sci Rep 7, 9741 (2017). 10.1038/s41598-017-09840-8

11 Huang, W. C., Ellis, J., Moody, P. C., Raven, E. L. & Roberts, G. C. Redox-linked domain movements in the catalytic cycle of cytochrome p_450_ reductase. Structure 21, 1581–1589 (2013). 10.1016/j.str.2013.06.022

12 Freeman, S. L. et al. Solution structure of the cytochrome P_450_ reductase-cytochrome. J Biol Chem 293, 5210–5219 (2018). 10.1074/jbc.RA118.001941

13 Zhang, L. et al. Structural insight into the electron transfer pathway of a self-sufficient P_450_ monooxygenase. Nat Commun 11, 2676 (2020). 10.1038/s41467-020-16500-5

14 Zhang, H. et al. The full-length cytochrome P_450_ enzyme CYP102A1 dimerizes at its reductase domains and has flexible heme domains for efficient catalysis. J Biol Chem 293, 7727–7736 (2018). 10.1074/jbc.RA117.000600

15 Askenasy, I. et al. Structure-Function Relationships in the Oligomeric NADPH-Dependent Assimilatory Sulfite Reductase. Biochemistry 57, 3764–3772 (2018). 10.1021/acs.biochem.8b00446

16 Siegel, L. M., Murphy, M. J. & Kamin, H. Reduced nicotinamide adenine dinucleotide phosphate-sulfite reductase of enterobacteria. I. The *Escherichia coli* hemoflavoprotein: molecular parameters and prosthetic groups. J Biol Chem 248, 251–264 (1973).

17 Murphy, M. J., Siegel, L. M., Tove, S. R. & Kamin, H. Siroheme: a new prosthetic group participating in six-electron reduction reactions catalyzed by both sulfite and nitrite reductases. Proc Natl Acad Sci U S A 71, 612–616 (1974).

18 Dailey, H. A. et al. Prokaryotic Heme Biosynthesis: Multiple Pathways to a Common Essential Product. Microbiol Mol Biol Rev 81 (2017). 10.1128/MMBR.00048-16

19 Nakayama, M., Akashi, T. & Hase, T. Plant sulfite reductase: molecular structure, catalytic function and interaction with ferredoxin. J Inorg Biochem 82, 27–32 (2000).

20 Schnell, R., Sandalova, T., Hellman, U., Lindqvist, Y. & Schneider, G. Siroheme- and [Fe4-S4]-dependent NirA from *Mycobacterium tuberculosis* is a sulfite reductase with a covalent Cys-Tyr bond in the active site. J Biol Chem 280, 27319–27328 (2005). https://doi.org/M502560200 [pii] 10.1074/jbc.M502560200

21 Pinto, R., Harrison, J. S., Hsu, T., Jacobs, W. R., Jr. & Leyh, T. S. Sulfite reduction in mycobacteria. J Bacteriol 189, 6714–6722 (2007). 10.1128/jb.00487-07

22 Jespersen, M., Pierik, A. J. & Wagner, T. Structures of the sulfite detoxifying F_420_-dependent enzyme from Methanococcales. Nat Chem Biol 19, 695–702 (2023). 10.1038/s41589-022-01232-y

23 Crane, B. R., Siegel, L. M. & Getzoff, E. D. Sulfite reductase structure at 1.6 A: evolution and catalysis for reduction of inorganic anions. Science 270, 59–67 (1995).

24 Crane, B. R., Siegel, L. M. & Getzoff, E. D. Probing the catalytic mechanism of sulfite reductase by X-ray crystallography: structures of the *Escherichia coli* hemoprotein in complex with substrates, inhibitors, intermediates, and products. Biochemistry 36, 12120–12137 (1997). 10.1021/bi971066i bi971066i [pii]

25 Crane, B. R., Siegel, L. M. & Getzoff, E. D. Structures of the siroheme- and Fe_4_S_4_-containing active center of sulfite reductase in different states of oxidation: heme activation via reduction-gated exogenous ligand exchange. Biochemistry 36, 12101–12119 (1997). 10.1021/bi971065q bi971065q [pii]

26 Crane, B. R. & Getzoff, E. D. The relationship between structure and function for the sulfite reductases. Curr Opin Struct Biol 6, 744–756 (1996). https://doi.org/S0959-440X(96)80003-0 [pii]

27 Young, L. J. & Siegel, L. M. Activated conformers of *Escherichia coli* sulfite reductase heme protein subunit. Biochemistry 27, 4991–4999 (1988).

28 Smith, K. W. & Stroupe, M. E. Mutational analysis of sulfite reductase hemoprotein reveals the mechanism for coordinated electron and proton transfer. Biochemistry 51, 9857–9868 (2012). 10.1021/bi300947a

29 Zeghouf, M., Fontecave, M. & Coves, J. A simplifed functional version of the *Escherichia coli* sulfite reductase. J Biol Chem 275, 37651–37656 (2000). 10.1074/jbc.M005619200 M005619200 [pii]

30 Murray, D. T., Weiss, K. L., Stanley, C. B., Nagy, G. & Stroupe, M. E. Small-angle neutron scattering solution structures of NADPH-dependent sulfite reductase. J Struct Biol 213, 107724 (2021). 10.1016/j.jsb.2021.107724

31 Tavolieri, A. M. et al. NADPH-dependent sulfite reductase flavoprotein adopts an extended conformation unique to this diflavin reductase. J Struct Biol (2019). 10.1016/j.jsb.2019.01.001

32 Walia, N. et al. Domain crossover in the reductase subunit of NADPH-dependent assimilatory sulfite reductase. J Struct Biol 215, 108028 (2023). 10.1016/j.jsb.2023.108028

33 Baba, T. et al. Construction of *Escherichia coli* K-12 in-frame, single-gene knockout mutants: the Keio collection. Mol Syst Biol 2, 2006 0008 (2006). msb4100050 [pii] 10.1038/msb4100050

34 Kampjut, D., Steiner, J. & Sazanov, L. A. Cryo-EM grid optimization for membrane proteins. iScience 24, 102139 (2021). 10.1016/j.isci.2021.102139

35 Levitz, T. S. et al. Approaches to Using the Chameleon: Robust, Automated, Fast-Plunge cryoEM Specimen Preparation. Front Mol Biosci 9, 903148 (2022). 10.3389/fmolb.2022.903148

36 Kim, J. Y., Nakayama, M., Toyota, H., Kurisu, G. & Hase, T. Structural and mutational studies of an electron transfer complex of maize sulfite reductase and ferredoxin. J Biochem (2016). 10.1093/jb/mvw016

37 Oliveira, T. F. et al. The crystal structure of *Desulfovibrio vulgaris* dissimilatory sulfite reductase bound to DsrC provides novel insights into the mechanism of sulfate respiration. J Biol Chem 283, 34141–34149 (2008). https://doi.org/M805643200 [pii] 10.1074/jbc.M805643200

38 Oliveira, T. F. et al. Structural insights into dissimilatory sulfite reductases: structure of desulforubidin from desulfomicrobium norvegicum. Front Microbiol 2, 71 (2011). 10.3389/fmicb.2011.00071

39 Parey, K., Warkentin, E., Kroneck, P. M. & Ermler, U. Reaction cycle of the dissimilatory sulfite reductase from *Archaeoglobus fulgidus*. Biochemistry 49, 8912–8921 (2010). 10.1021/bi100781f

40 Schiffer, A. et al. Structure of the dissimilatory sulfite reductase from the hyperthermophilic archaeon *Archaeoglobus fulgidus*. J Mol Biol 379, 1063–1074 (2008). https://doi.org/S0022-2836(08)00456-7 [pii] 10.1016/j.jmb.2008.04.027

41 Gruez, A. et al. Four crystal structures of the 60 kDa flavoprotein monomer of the sulfite reductase indicate a disordered flavodoxin-like module. J Mol Biol 299, 199–212 (2000). 10.1006/jmbi.2000.3748 S0022-2836(00)93748-3 [pii]

42 Zhong, E. D., Bepler, T., Berger, B. & Davis, J. H. CryoDRGN: reconstruction of heterogeneous cryo-EM structures using neural networks. Nat Methods 18, 176–185 (2021). 10.1038/s41592-020-01049-4

43 Xia, C. et al. Conformational changes of NADPH-cytochrome P_450_ oxidoreductase are essential for catalysis and cofactor binding. J Biol Chem 286, 16246–16260 (2011). 10.1074/jbc.M111.230532

44 Flocco, M. M. & Mowbray, S. L. Planar stacking interactions of arginine and aromatic side-chains in proteins. J Mol Biol 235, 709–717 (1994). 10.1006/jmbi.1994.1022

45 D’Imprima, E. et al. Protein denaturation at the air-water interface and how to prevent it. Elife 8 (2019). 10.7554/eLife.42747

46 Glaeser, R. M. et al. Factors that Influence the Formation and Stability of Thin, Cryo-EM Specimens. Biophys J 110, 749–755 (2016). 10.1016/j.bpj.2015.07.050

47 Glaeser, R. M. & Han, B. G. Opinion: hazards faced by macromolecules when confined to thin aqueous films. Biophys Rep 3, 1–7 (2017). 10.1007/s41048-016-0026-3

48 Glaeser, R. M. PROTEINS, INTERFACES, AND CRYO-EM GRIDS. Curr Opin Colloid Interface Sci 34, 1–8 (2018). 10.1016/j.cocis.2017.12.009

49 Wu, J. Y., Siegel, L. M. & Kredich, N. M. High-level expression of *Escherichia coli* NADPH-sulfite reductase: requirement for a cloned *cysG* plasmid to overcome limiting siroheme cofactor. J Bacteriol 173, 325–333 (1991).

50 Suloway, C. et al. Automated molecular microscopy: the new Leginon system. J Struct Biol 151, 41–60 (2005). S1047-8477(05)00072-9 [pii] 10.1016/j.jsb.2005.03.010

51 Suloway, C. et al. Automated molecular microscopy: The new Leginon system. Journal of Structural Biology 151, 41–60 (2005). 10.1016/j.jsb.2005.03.010

52 Cheng, A. et al. Leginon: New features and applications. Protein Sci 30, 136–150 (2021). 10.1002/pro.3967

53 Zheng, S. Q. et al. MotionCor2: anisotropic correction of beam-induced motion for improved cryo-electron microscopy. Nat Methods 14, 331–332 (2017). 10.1038/nmeth.4193

54 Scheres, S. H. RELION: implementation of a Bayesian approach to cryo-EM structure determination. J Struct Biol 180, 519–530 (2012). 10.1016/j.jsb.2012.09.006

55 Rohou, A. & Grigorieff, N. CTFFIND4: Fast and accurate defocus estimation from electron micrographs. J Struct Biol 192, 216–221 (2015).

56 Punjani, A., Rubinstein, J. L., Fleet, D. J. & Brubaker, M. A. cryoSPARC: algorithms for rapid unsupervised cryo-EM structure determination. Nat Methods 14, 290–296 (2017). 10.1038/nmeth.4169

57 Aiyer, S., Zhang, C., Baldwin, P. R. & Lyumkis, D. Evaluating Local and Directional Resolution of Cryo-EM Density Maps. Methods Mol Biol 2215, 161–187 (2021). 10.1007/978-1-0716-0966-8_8

58 Punjani, A., Zhang, H. & Fleet, D. J. Non-uniform refinement: adaptive regularization improves single-particle cryo-EM reconstruction. Nat Methods 17, 1214–1221 (2020). 10.1038/s41592-020-00990-8

59 Sanchez-Garcia, R. et al. DeepEMhancer: a deep learning solution for cryo-EM volume post-processing. Commun Biol 4, 874 (2021). 10.1038/s42003-021-02399-1

60 Bepler, T. et al. Positive-unlabeled convolutional neural networks for particle picking in cryo-electron micrographs. Nat Methods 16, 1153–1160 (2019). 10.1038/s41592-019-0575-8

61 Punjani, A. & Fleet, D. 3D Flexible Refinement: Determining Structure and Motion of Flexible Proteins from Cryo-EM. Microsc Microanal 29, 1024 (2023). 10.1093/micmic/ozad067.518

62 Pettersen, E. F. et al. UCSF Chimera - A visualization system for exploratory research and analysis. Journal of Computational Chemistry 25, 1605–1612 (2004). 10.1002/jcc.20084

63 Jumper, J. et al. Highly accurate protein structure prediction with AlphaFold. Nature 596, 583–589 (2021). 10.1038/s41586-021-03819-2

64 Roney, J. P. & Ovchinnikov, S. State-of-the-Art Estimation of Protein Model Accuracy Using AlphaFold. Phys Rev Lett 129, 238101 (2022). 10.1103/PhysRevLett.129.238101

65 Adams, P. D. et al. PHENIX: a comprehensive Python-based system for macromolecular structure solution. Acta Crystallogr D Biol Crystallogr 66, 213–221 (2010). https://doi.org/S0907444909052925 [pii] 10.1107/S0907444909052925

66 Emsley, P., Lohkamp, B., Scott, W. G. & Cowtan, K. Features and development of Coot. Acta Crystallogr D Biol Crystallogr 66, 486–501 (2010). https://doi.org/S0907444910007493 [pii] 10.1107/S0907444910007493

