## Supplemental material for "Structure of dimerized assimilatory NADPH-dependent sulfite reductase reveals the minimal interface for diflavin reductase binding"

**Corresponding Author:** M. Elizabeth Stroupe

| Variable | SiRFP-43/SiRHP | SiRFP-60Δ/SiRHP | SiRFP-60X/SiRHP |
|---|---|---|---|
| **Expression** | co-lysis | co-lysis | reconstitution |
| **SiRFP variant (including the amino acids expressed) and the mass from which the name is derived** | N-terminal octamerization and Fld- domain truncated, monomeric 43 kDa FNR domain (amino acids 237-599) | 60 kDa monomer with linker truncation (ΔAAPSQS) (amino acids 53-599 minus 212-217) | 60 kDa crosslinked monomer with four engineered variants: C162T, C552S, E121C, and N556C (amino acids 53-599) |
| **Theoretical mass of the complex (kDa)** | 107 | 123 | 124 |
| **Ability to complement SiRFP-deficient *Escherichia coli* without exogenous $S^{2-}$** | - | - | - |

**Table S1**. Expression system, nomenclature, mass, and activity for each heterodimer.

| Sample assembly | SiRFP-43/SiRHP | SiRFP-60Δ/SiRHP | SiRFP-60X/SiRHP |
|---|---|---|---|
| **SiRFP-43/SiRHP** | Titan Krios | Titan Krios | Titan Krios |
| **Voltage (KV)** | 300 | 300 | 300 |
| **Camera** | K3 | K3 | Apollo |
| **Magnification** | 105 K | 105 K | 59 K |
| **Total dose (e/Å²)** | 60 | 60 | 60 |
| **Pixel size (Å)** | 0.844 | 0.844 | 0.765 |
| **Number of movies** | 14,628 | 25,488 | 9,963 |
| **Number of particles** | 1,500,000 | 550,000 | 179,000 |
| **Resolution (Å)** | 3.47* | 3.49 | 2.78 |
| **Data collection location** | NYSBC | NYSBC | FSU BSIR |

**Table S2.** Data collection parameters and final reconstruction information.
*Nominal resolution

| Parameter | Value |
|---|---|
| **Mean B-factor (Å²)** | 57.2 |
| **Map CC** | 0.86 |
| **RMSD [bonds] (Å)** | 0.003 |
| **RMSD [angles] (Å)** | 0.632 |
| **All-atom clash-score** | 7.3 |
| **Ramachandran plot** | |
| Favored (%) | 96.4 |
| Allowed (%) | 3.50 |
| Outliers (%) | 0.09 |
| **Rotamer outliers (%)** | 0.98 |
| **C-β deviations (%)** | 0.00 |
| **PDB ID** | 9C91 |
| **EMDB ID** | EMD-45359 |

**Table S3.** Model refinement and validation statistics.

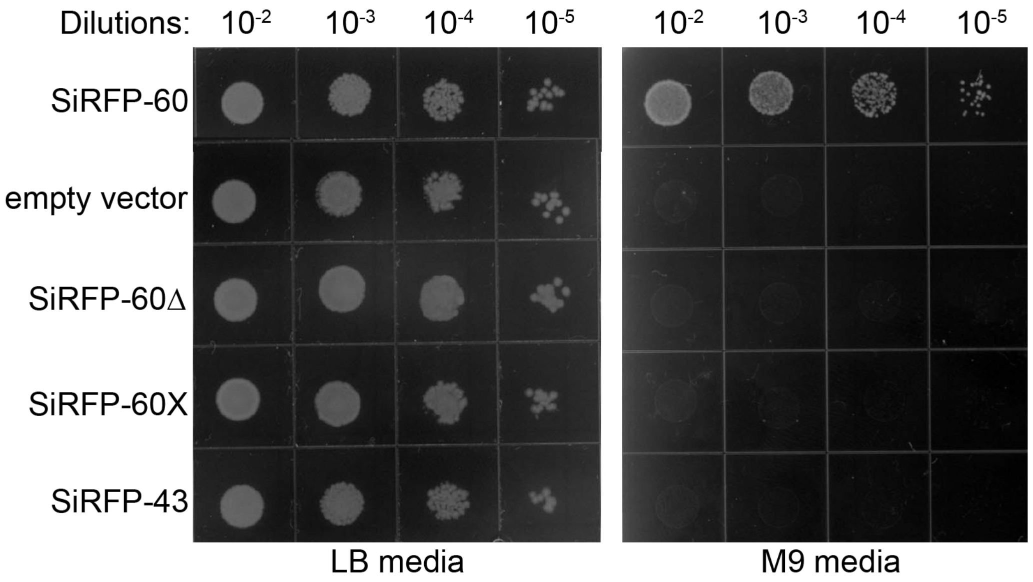


**Fig S1.** The SiRFP variants studied here do not complement SiRFP-deficient *E. coli* under reduced-sulfur growth conditions.

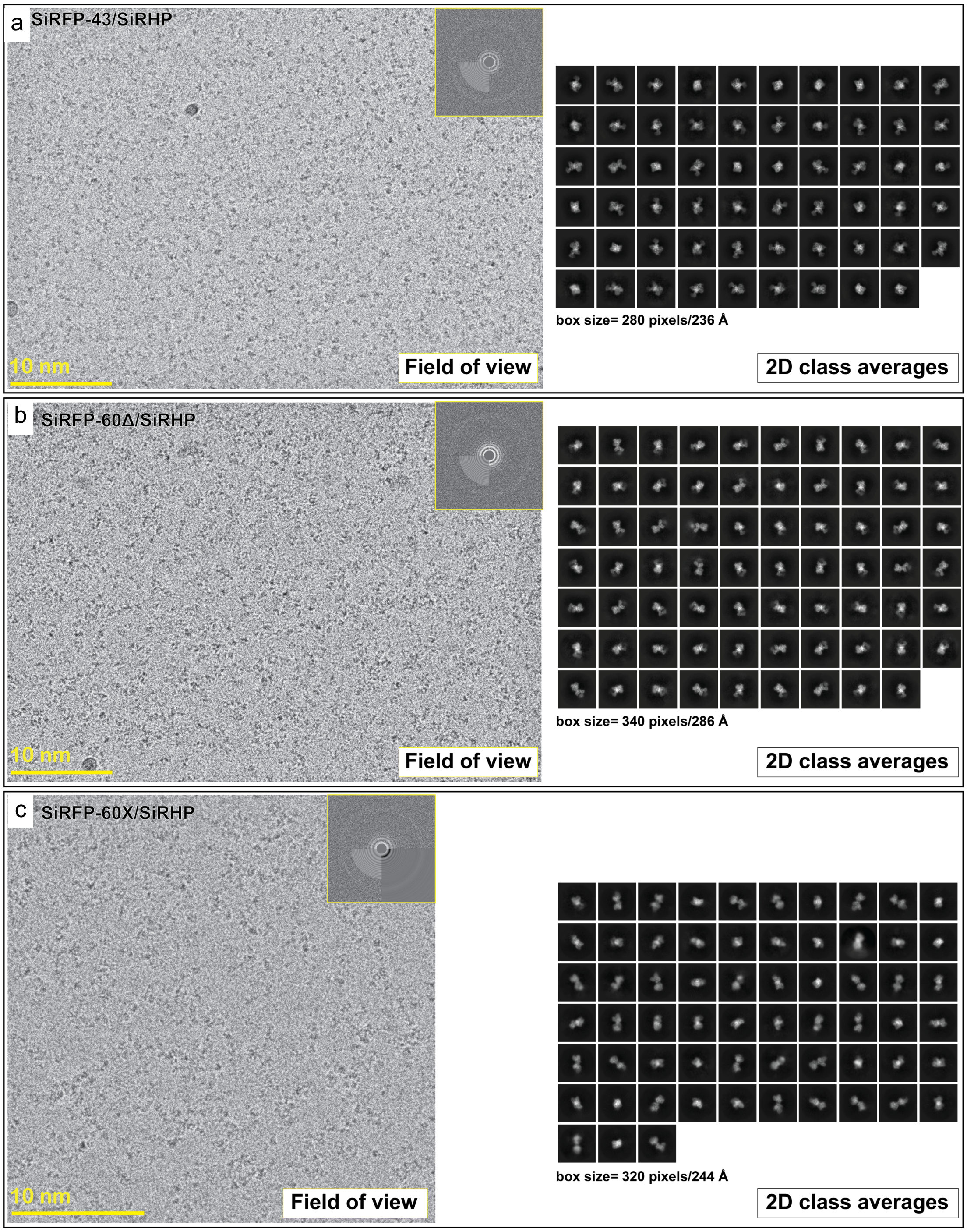


**Fig S2.** Raw data for each SiR dimer **a** Exemplar micrograph and selected 2D class averages from SiRFP-43/SiRHP dataset. **b** Exemplar micrograph and selected 2D class averages from SiRFP-60Δ/SiRHP dataset. **c** Exemplar micrograph and selected 2D class averages from SiRFP-60X/SiRHP dataset.

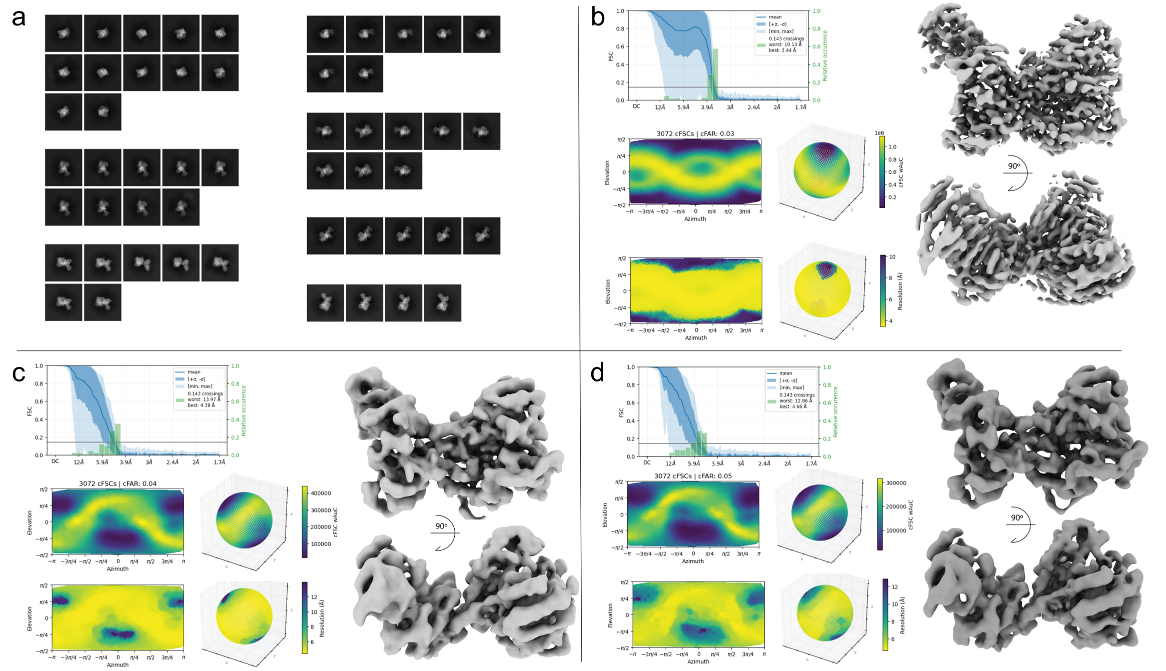


**Fig S3.** Orientation analysis of the SiRFP-43/SiRHP dimer **a** 2D superclass averages from the SiRFP-43/SiRHP dataset used to reduce the particles in each cluster to block preferred orientation effect. **b** Non-uniform refinement and orientation diagnostics results for all particles with good 2D class averages features (~1,500,000 particles). **c** Non-uniform refinement and orientation diagnostics results for reduced number of particles based on 2D rebalance super classification 350,000 particles). **d** Non-uniform refinement and orientation diagnostics results for reduced number of particles based on 2D rebalance super classification 105,000 particles).

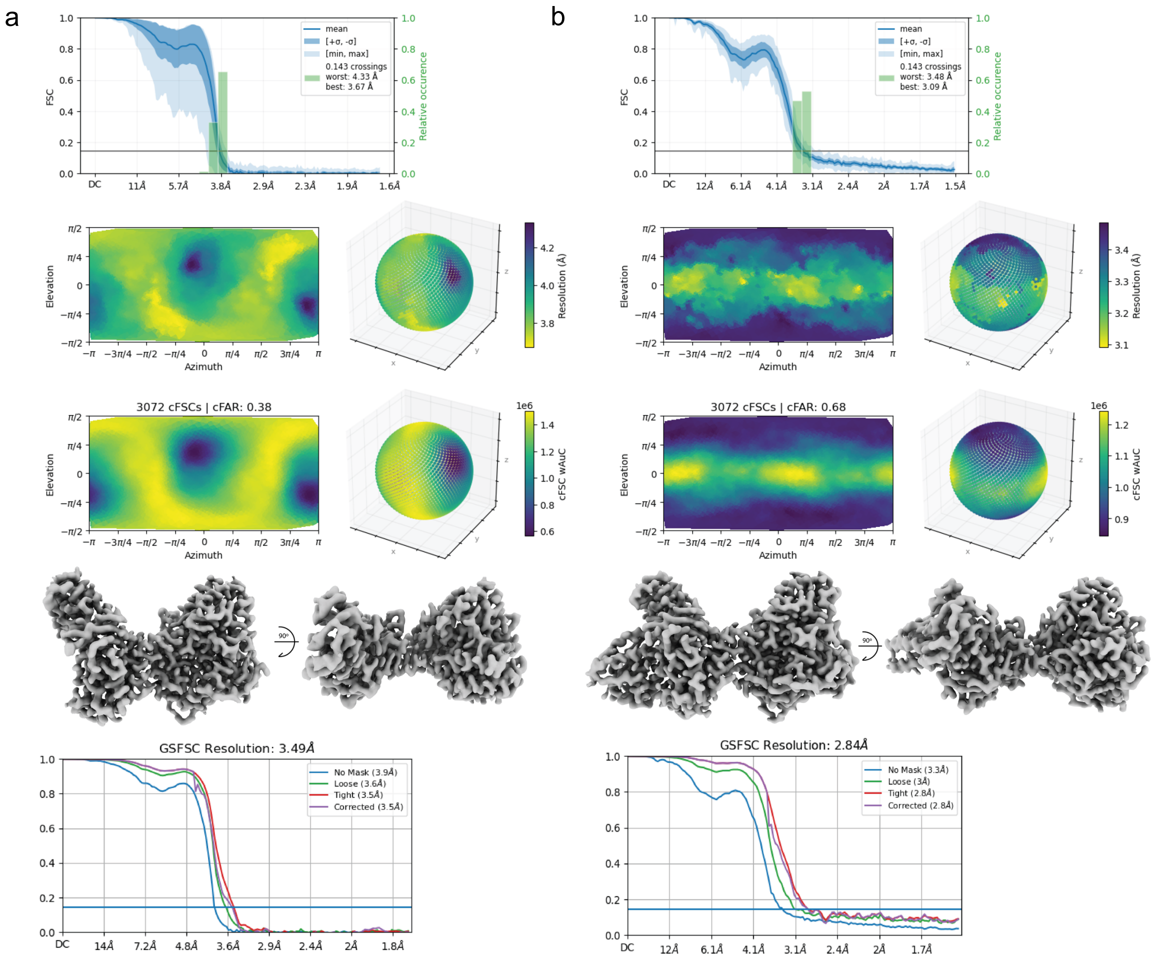


**Fig S4.** 3D orientation analysis for SiRFP-60Δ/SiRHP and SiRFP-60X/SiRHP

**a** Orientation diagnostics results, highest resolution map and the refinement's GSFSC curve for SiRFP-60Δ/SiRHP particles.

**b** Orientation diagnostics results, highest resolution map and the refinement's GSFSC curve for SiRFP-60X/SiRHP particles.

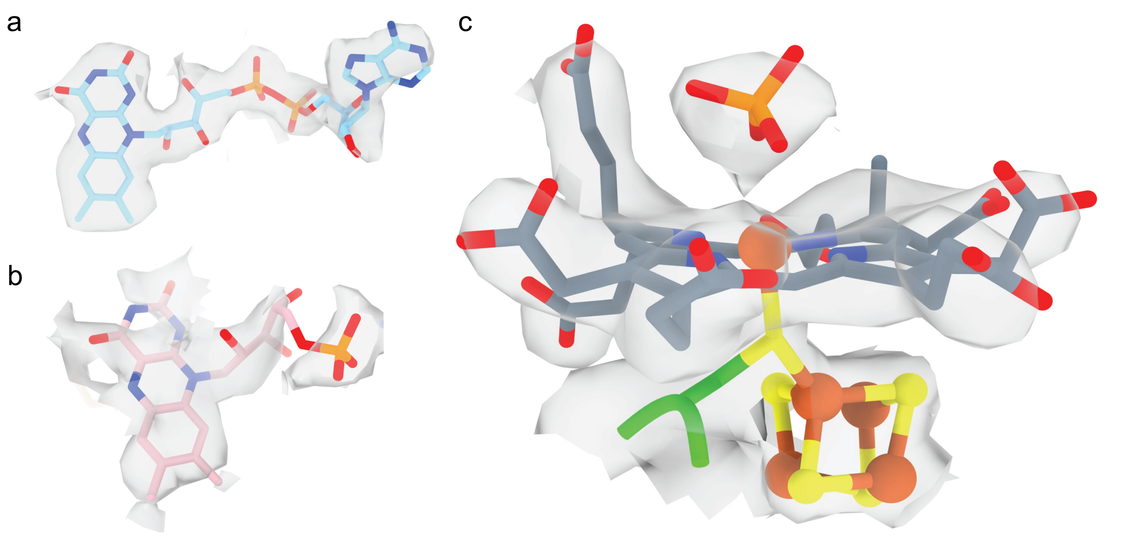


**Fig S5.** SiRFP-60X/SiRHP high resolution map (2.78 Å) with placed co-enzymes: **a** FAD (contour level 0.5), **b** FMN (contour level 0.3), **c** Siroheme-$Fe_4S_4$ cluster bound to a distal phosphate with the bridging h-Cys483 (contour level 0.8).

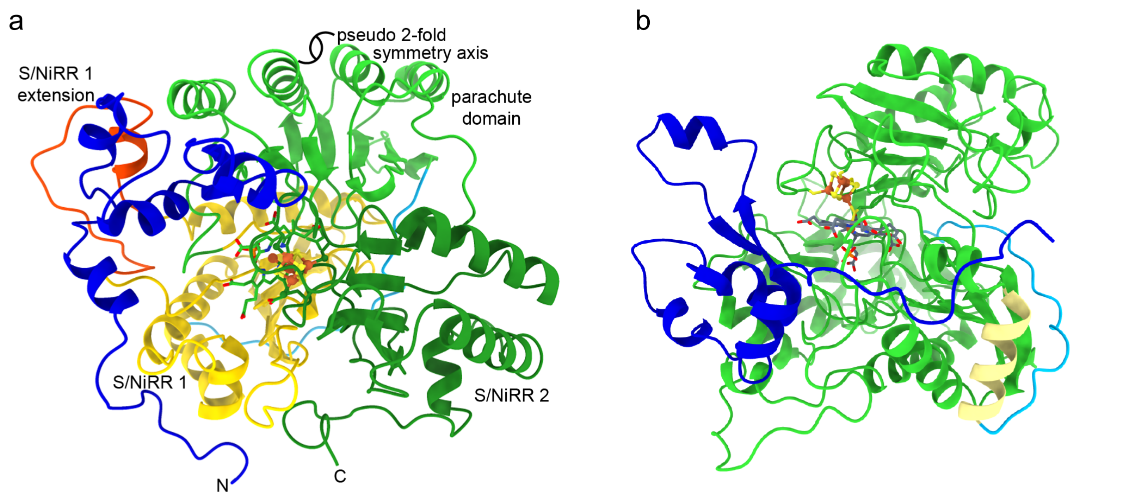


**Fig S6.** Path of SiRHP's previously undetermined N-terminal 80 amino acids. **a** The N-terminus of SiRHP (blue) breaks the pseudo 2-fold symmetry that relates S/NiRR 1 (domain 1, yellow + the N-terminus of the parachute domain, light green) and S/NiRR 2 (domain 2, dark green + the C-terminus of the parachute domain, light green. The S/NiRR 1 extension is orange. **b** Blue: amino acids 1-80. Green: S/NiRR 1 and 2. Yellow: helix preceding the siroheme mimic (h11). Cyan: siroheme mimic joining S/NiRRs 1 and 2. **b** is rotated $90^o$ to the right and $90^o$ into the place of the page relative to the view in **a**.

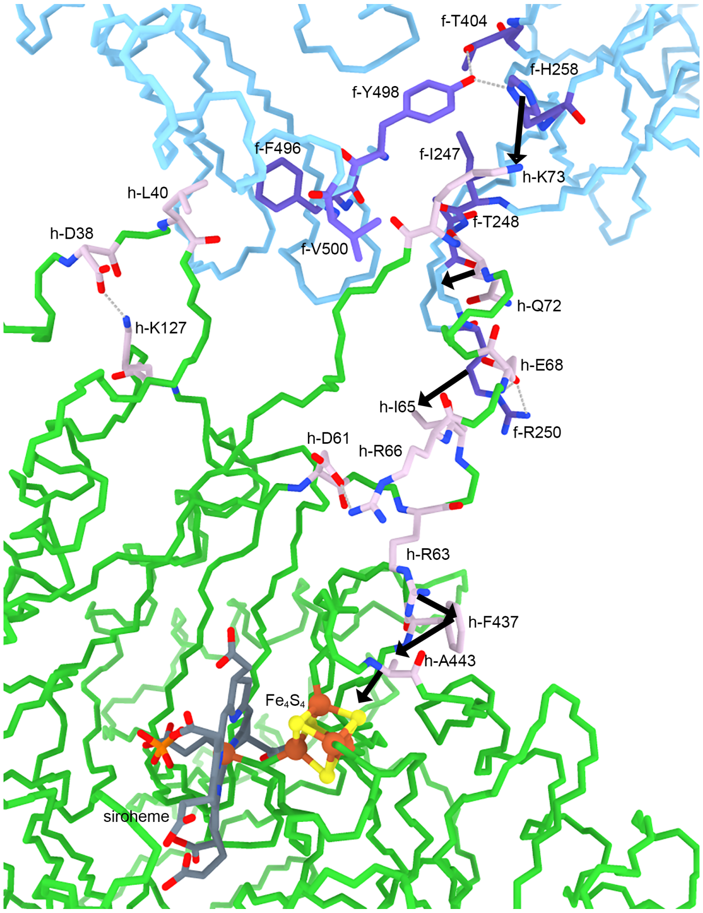


**Fig S7.** An extensive network of hydrophobic interactions, ionic bonds, and hydrogen bonds bridge from the SiRFP/SiRHP interface to the active site. Through space ionic or hydrogen bonds are represented by gray dashed lines. Through space hydrophobic interactionsare represented by black arrows.

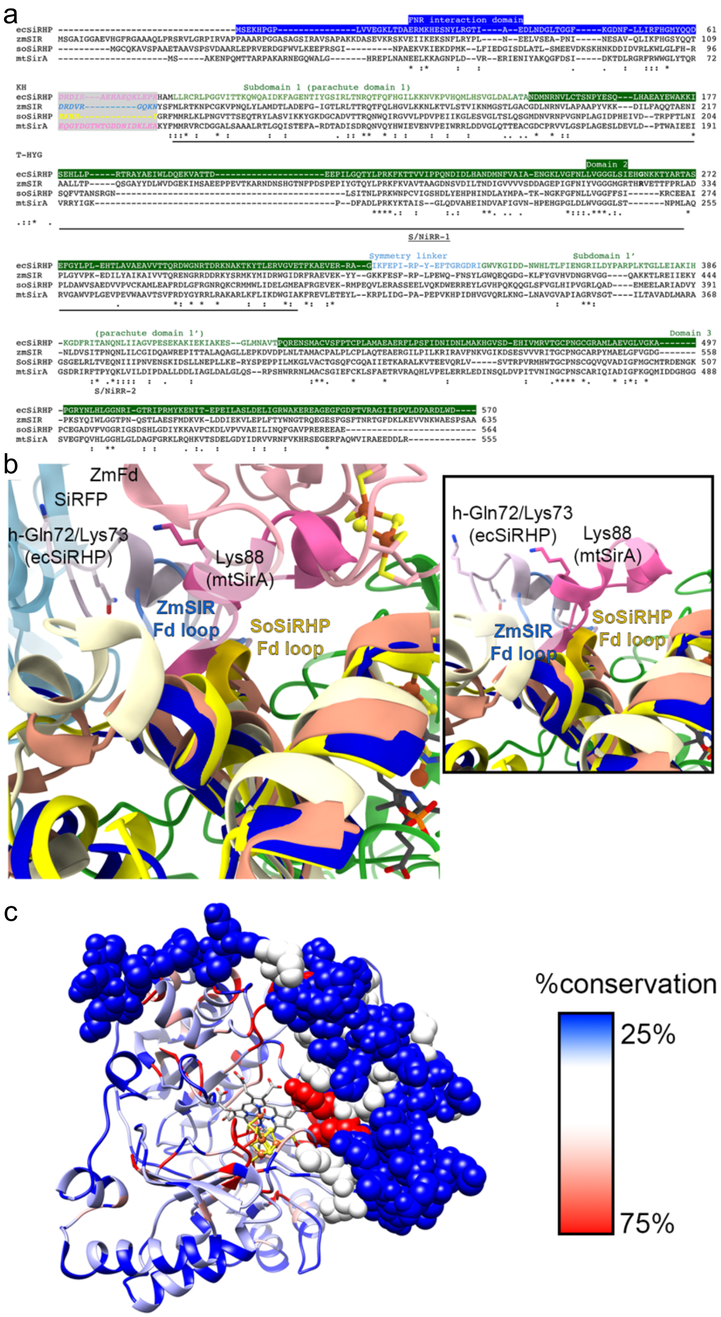


**Fig. S8.** The loop that contains h-Lys73 differs in both **a** sequence (structure-based sequence alignment in UCSF Chimera[1]) and **b** structure, amongst diflavin-reductase (sky blue) dependent hemoprotein (like in *E. coli*, mauve) and ferredoxin (light pink) dependent (like in *Z. mays*, cornflower blue, *S. oleracea*, dark yellow, or *M. tuberculosis*, dark pink). ZmFd and SiRFP have been removed in the inset for clarity. N-terminus of ecSiRHP is dark blue, zmSIR is yellow, soSiRHP is light yellow, and mtSir is peach. **c** The N-terminus of SiRHP is not strongly conserved in comparison to the active site amino acids, as calculated in UCSF Chimera[1].

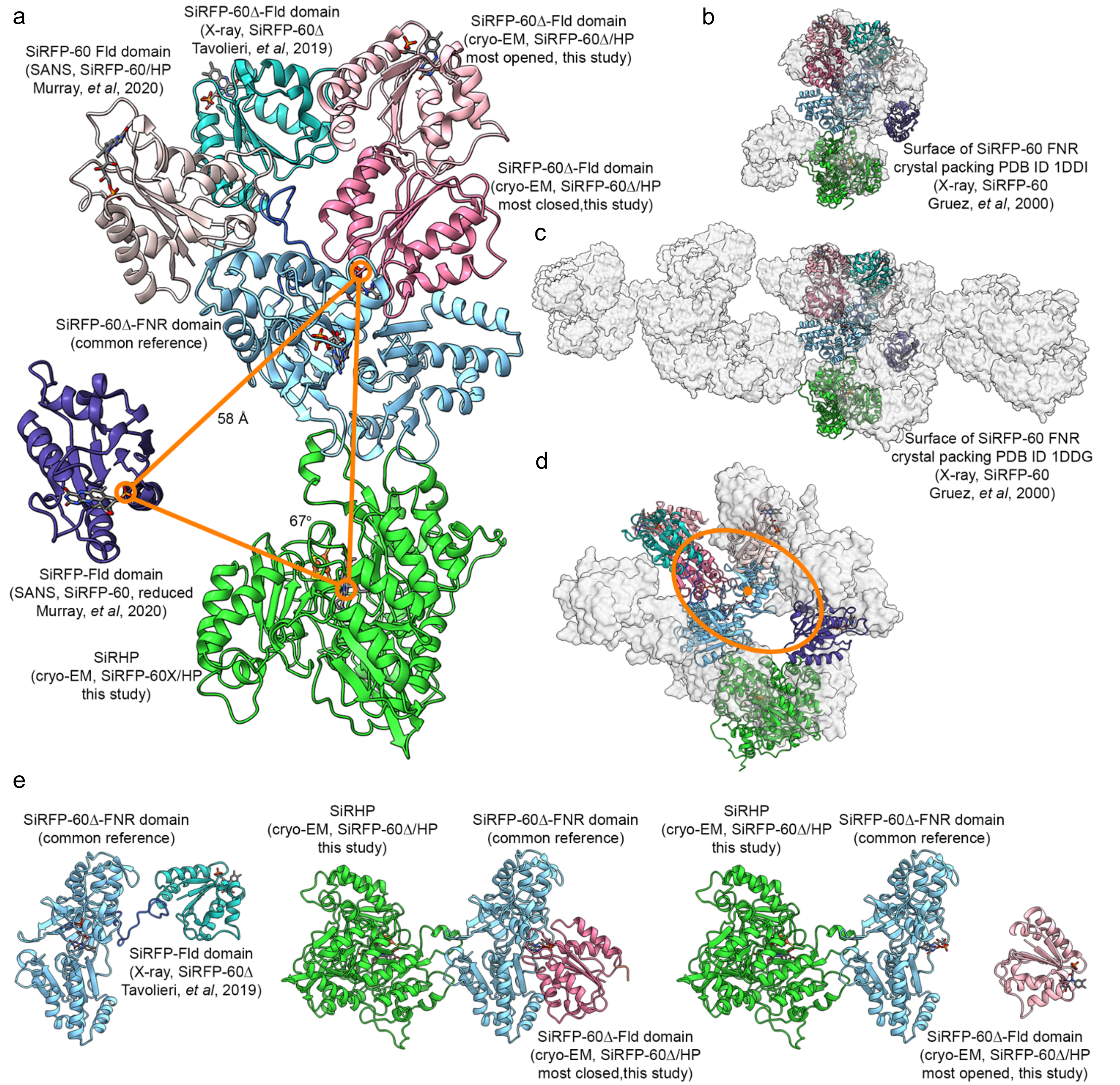


**Fig. S9.** The SiRFP-60 Fld domain is mobile. **a** Each position was determined by cryo-EM (this study, showing the most extreme positions from the SiRFP-60Δ/SiRHP series of structures), X-ray crystallography[2], or SANS[3]. The position of the Fld domain is dependent on the SiRFP oxidation state or SiRHP binding. **b** When SiRFP-60 is crystallized, the Fld domain is not ordered although it is present[4], leaving large solvent channels that can accommodate the positions of the Fld domain captured by SANS[3], X-ray crystallography of SiRFP-60Δ[2], or cryo-EM of SiRFP-60Δ/SiRHP. **c** Another SiRFP-60 crystal form[4] has solvent channels that accommodate the Fld domain in some positions. **b** and **c** are oriented 90° to the left around a vertical axis from where they are positioned in **a** and colored the same. SiRHP is only present in the cryo-EM structure. **d** The Fld domain can adopt positions along an elliptical cone shape relative to the FNR domain, accommodated in the channels from the crystal form shown in **b** (rotated 90° to the left and out of the page relative to **b**). **e** SiRFP-60Δ is shown with the FNR domain in a common position to demonstrate the different orientations of the Fld domain. The linker is not visible in this cryo-EM reconstruction. The most extreme closed or open positions of the Fld domain are shown. The closed position is similar to that seen in SiRFP-60X/SiRHP.

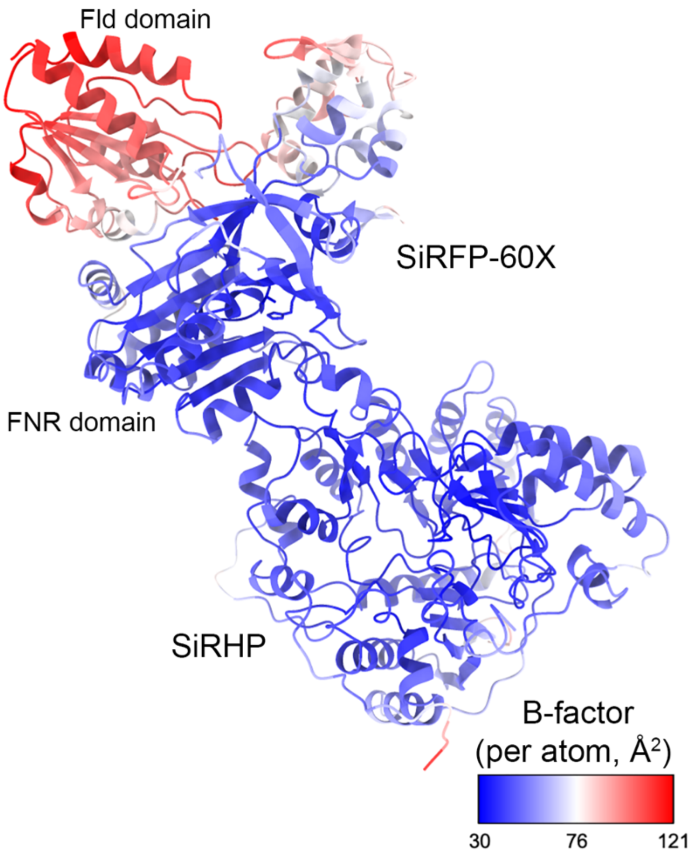


**Fig. S10** The Fld domain in SiRFP has the highest B-factors in the SiRFP-60X/SiRHP dimer.

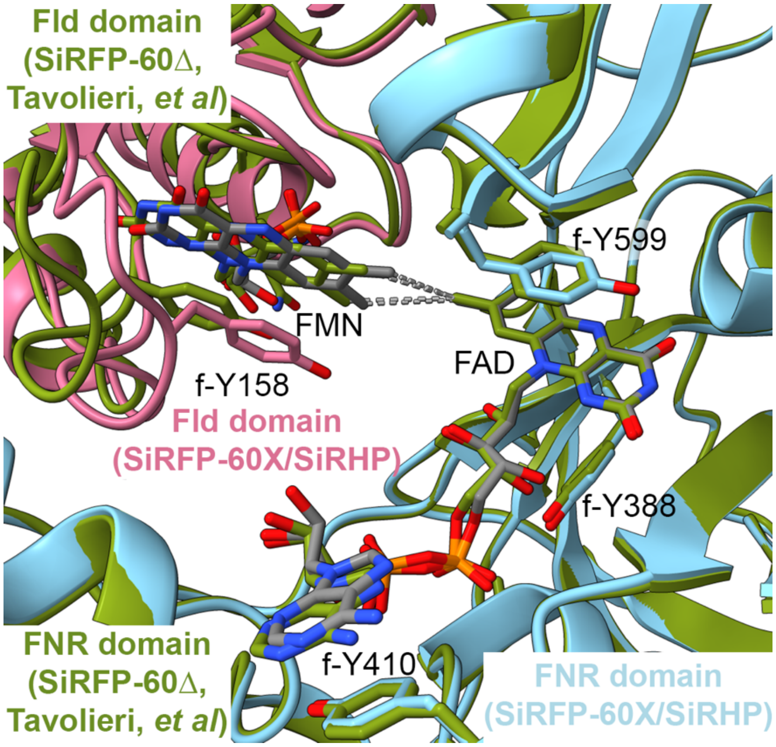


**Fig. S11.** SiRHP binding does not effect flavin (gray) binding to SiRFP-60X (pink and light blue) compared to their binding in SiRFP-60Δ[2] (olive green). The crosslink between the Fld (pink) and FNR (blue) domains shifts the FMN and the FMN binding loop towards the FNR domain.

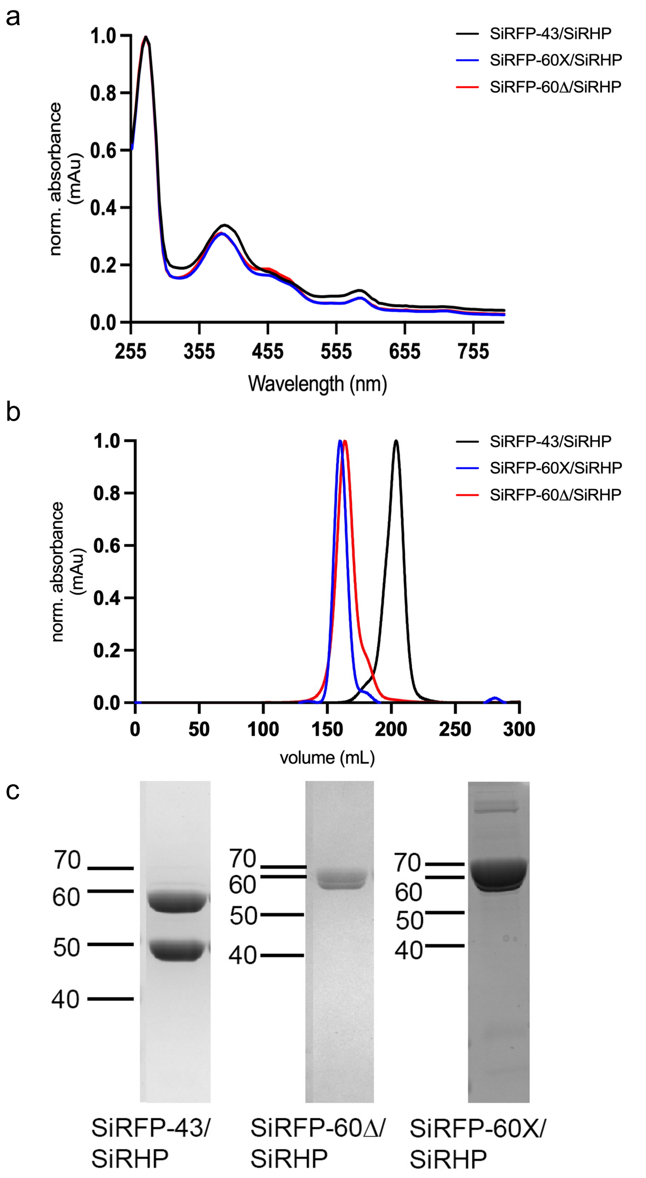


**Fig S12.** Biochemical analysis of each SiRFP/SiRHP variant shows that all are properly formed. **a** UV-Visible spectra showed that all variants are properly folded with cofactors bound. **b** Size exclusion chromatography profiles of all three variants with their characteristic elution profiles. **c** SDS-PAGE analysis of the variants, each of the protein preparation studied here, showed the expected molecular weights (kDa) for the subunits of the purified complexes.

**Video S1.** Swinging Fld domain shown in motion, obtained from SiRFP-60Δ/SiRHP dataset via CryoDRGN showing one of the latent spaces out of two.

### REFERENCES


1 Pettersen, E. F. *et al.* UCSF Chimera - A visualization system for exploratory research and analysis. *Journal of Computational Chemistry* **25**, 1605-1612 (2004). https://doi.org/10.1002/jcc.20084

2 Tavolieri, A. M. *et al.* NADPH-dependent sulfite reductase flavoprotein adopts an extended conformation unique to this diflavin reductase. *J Struct Biol* (2019). https://doi.org/10.1016/j.jsb.2019.01.001

3 Murray, D. T., Weiss, K. L., Stanley, C. B., Nagy, G. & Stroupe, M. E. Small-angle neutron scattering solution structures of NADPH-dependent sulfite reductase. *J Struct Biol* **213**, 107724 (2021). https://doi.org/10.1016/j.jsb.2021.107724

4 Gruez, A. *et al.* Four crystal structures of the 60 kDa flavoprotein monomer of the sulfite reductase indicate a disordered flavodoxin-like module. *J Mol Biol* **299**, 199-212 (2000). https://doi.org/10.1006/jmbi.2000.3748 S0022-2836(00)93748-3 [pii]
